# Rapid and reversible regulation of cell cycle progression in budding yeast using optogenetics

**DOI:** 10.1101/2024.09.21.614242

**Authors:** Andriana Koutsoumpa, Andreas Milias-Argeitis

## Abstract

The regulatory complexity of the eukaryotic cell cycle poses technical challenges in experiment design and data interpretation, leaving gaps in our understanding of how cells coordinate cell cycle-related processes. Traditional methods, such as knockouts and deletions are often ineffective to compensatory interactions in the cell cycle control network, while chemical agents that cause cell cycle arrest can have undesired pleiotropic effects. Synthetic inducible systems targeting specific cell cycle regulators offer potential solutions but are limited by the need for external inducers, which make fast reversibility technically challenging. To address these issues, we developed an optogenetic tool (OPTO-Cln2) that enables light-controlled and reversible regulation of G1 progression in budding yeast. Through extensive validation and benchmarking via time-lapse microscopy, we verify that OPTO-Cln2-carrying strains can rapidly toggle between normal and altered G1 progression. By integrating OPTO-Cln2 with a readout of nutrient-sensing pathways (TORC1 and PKA), we show that the oscillatory activity of these pathways is tightly coordinated with G1 progression. Finally, we demonstrate that the rapid reversibility of OPTO-Cln2 facilitates multiple cycles of synchronous arrest and release of liquid cell cultures. Our work provides a powerful new approach for studying cell cycle dynamics and the coordination of growth- with division-related processes.

## Introduction

The eukaryotic cell cycle comprises a series of processes through which cells grow, duplicate their DNA and divide (Morgan, 2007). These processes are regulated in space and time through the coordinated action of signaling systems, transcriptional networks and metabolic pathways. Despite significant progress in the study of cell cycle control in the past decades, the complex feedback interactions within and among these regulatory mechanisms, along with multiple levels of redundancy, continue to present technical challenges in experiment design and data interpretation. Consequently, our understanding of the dynamic interplay of the various cell cycle regulators and the coordination of growth- with division-related processes remains incomplete (Øvrebø *et al*, 2022; Liu *et al*, 2022b, 2022a; Fisher & Krasinska, 2022), even in unicellular eukaryotes like budding yeast (*Saccharomyces cerevisiae*) (Brambila *et al*, 2024; Garmendia-Torres *et al*, 2018; Sommer *et al*, 2021).

Tools that induce perturbations to physiological cell cycle progression have been indispensable for the study of the cell cycle control system. Deletions and mutations of key cell cycle regulators (e.g. temperature-sensitive mutants) and chemical agents that induce cell cycle arrest are some of the earliest tools used to alter cell cycle progression (Noguchi & Gadaleta, 2014). While invaluable for early cell cycle studies, these tools have several drawbacks (Banfalvi, 2017). The effects of deletions and mutations can be masked by redundant pathways and compensatory feedback loops; blocking agents can induce unwanted stress responses and affect cell viability; and temperature-sensitive mutants are often hypoactive at the permissive temperature and exhibit incomplete loss of function at non-permissive temperatures. Additionally, these mutants require shifting organisms to non-physiological temperatures, which may in turn induce stress responses.

The advent of synthetic biology techniques has enabled the application of conditional knockout (Yesbolatova *et al*, 2020) and overexpression (Gligorovski *et al*, 2020; Ottoz *et al*, 2014) systems to target specific cell cycle components. Alongside the development of chemicals that specifically target cell cycle regulators (Łukasik *et al*, 2021; Bavetsias & Linardopoulos, 2015; Liu, 2015), a diverse set of tools is now available for inducing targeted and dynamic perturbations to cell cycle progression. However, a key limitation of these systems is fast reversibility: even when gene expression or protein destabilization can be switched reversibly, this operation typically requires the addition and removal of small molecule inducers from the culture medium. The same principle applies to molecules that selectively modulate the activity of cell cycle components. Dynamic manipulation of molecule concentrations often requires complex and expensive equipment such as microfluidic platforms and perfusion systems, and can be challenging to scale up for large cell cultures.

Optogenetics, a technique that enables fast and targeted light-based modulation of protein activity (Repina *et al*, 2017), can be used to reversibly regulate the activity of specific cell cycle components with high temporal resolution, thereby overcoming the challenges mentioned above. However, the application of optogenetics in cell cycle research has remained relatively unexplored. To address the lack of tools for light-based cell cycle control, we developed and characterized a transcription-based optogenetic system for the regulation of G1 progression in budding yeast.

While deletion of all three G1 cyclins of yeast results in cell cycle arrest (Richardson *et al*, 1989), cells with a single G1 cyclin are viable and exhibit a prolonged G1 phase. Thus, it is possible to modulate G1 length or completely halt the cell cycle by controlling the expression of a G1 cyclin in a background strain lacking one or two of the other G1 cyclins. To reversibly prolong or halt G1 progression, we developed the OPTO-Cln2 system. OPTO-Cln2 leverages the light-driven changes in the activity of the EL222 transcription factor (Motta-Mena *et al*, 2014) to regulate the expression of a G1 cyclin Cln2 from an EL222-responsive promoter. This promoter can be tightly switched off in darkness and achieves physiological Cln2 levels under low-intensity blue light. When introduced in a strain lacking another the G1 cyclin Cln3, OPTO-Cln2 enabled reversible switching between the wild-type and *cln2Δcln3Δ* phenotypes, effectively toggling between fast and slow G1 progression. On the other hand, introduction of OPTO-Cln2 in a background deleted for the G1 cyclins Cln1 and Cln3 allowed us to reversibly arrest and release the cell cycle.

To verify that our optogenetic strains functioned as expected, we used time-lapse microscopy to characterize their phenotypes and switching dynamics. Additionally, we compared the cell cycle progression and morphology of these strains to the corresponding features of two G1 cyclin deletion strains, one of which had remained almost fully uncharacterized. Our detailed observations uncovered cellular behaviors that cannot be fully explained by the established model of G1 regulation and suggest the presence of compensatory mechanisms that alter the levels or timing of G1 cyclin expression in deletion mutants, possibly in connection with an increase in cell size. Prompted by these observations, we used a previously established single-cell readout (Guerra *et al*, 2022) to monitor the cell cycle dynamics of two major and highly conserved nutrient-sensing pathways (TORC1 and PKA) that regulate cell growth. We observed that the joint TORC1 and PKA activity shows a maximum during G1 and minima at mitosis and budding (entry to S) under both light and dark conditions in our optogenetic strains, suggesting that TORC1 and PKA activity during G1 is tightly coordinated with G1 progression. On the other hand, uncoupling growth from division in G1-arrested cells led to persistently high signaling activity, supporting the notion that entry to the S phase is accompanied by an attenuation of TORC1 and PKA activity.

As a final demonstration of the capabilities of our optogenetic system, we monitored the population dynamics of cultures subjected to light-induced cell cycle arrest and release. Our results showed that the OPTO-Cln2 system achieves a performance very similar to that of alpha factor treatment (Breeden, 1997; Amon, 2002), the current gold standard arrest-and-release method for budding yeast. Contrary to that method, however, our approach does not require the addition and removal of inducers, and therefore enables multiple rounds of arrest and release in large culture volumes.

Overall, our work demonstrates that the advent of optogenetics for cell cycle regulation holds the potential to transform our understanding of cell cycle dynamics and open new possibilities for biotechnological applications. The conserved nature of G1 progression mechanisms suggests that our approach can be extrapolated to higher eukaryotic organisms, facilitating cell cycle studies of medically important cell types such as stem cells and tumor cells.

## Results

### Deletion of G1 cyclins increases G1 duration in daughter cells and causes an increase in cell size

Since our optogenetic approach to cell cycle control is centered around the regulation of G1 progression, we first sought to establish the conditions under which this regulation can be effective and the phenotypes than can be attained under different light conditions. For this purpose, we characterized the proliferation of yeast cells carrying G1 cyclin deletions.

Budding yeast cells missing one or two G1 cyclins are still able to progress through G1, whereas cells missing all three G1 cyclins arrest at the G1/S boundary (Richardson *et al*, 1989). To determine how G1 cyclin deletion mutants differ from the prototrophic YSBN6 reference strain (Canelas *et al*, 2010) (henceforth abbreviated as “WT”) in terms of cell cycle progression and morphology, we constructed strains lacking one (*cln3Δ*), or two (*cln2Δcln3Δ*) G1 cyclins in the YSBN6 background (Materials and Methods).

Estimation of population doubling times via flow cytometry showed that the *cln2Δcln3Δ* mutant proliferated only slightly slower than the WT (Supp. Fig. S1). To observe in more detail how cell cycle progression and cell size are affected by G1 cyclin deletions, we used time-lapse microscopy to determine the distributions of G1 and post-G1 phase durations (roughly corresponding to the unbudded vs. budded part of the cell cycle, cf. Methods), as well as the distributions of cell volumes at key cell cycle events (Methods) in mother and daughter cells. The distinction between mothers and daughters was necessary because of their different size requirements for cell cycle commitment (Rupeš, 2002), which lead to differences in cell cycle progression (Fig. 1B). Full distributions of our measurement are provided in Fig. 1, while Supplementary Tables 5-8 summarize the distribution medians (which are also used throughout in the text) and provide statistical comparisons between different strains.

**Figure 1.**
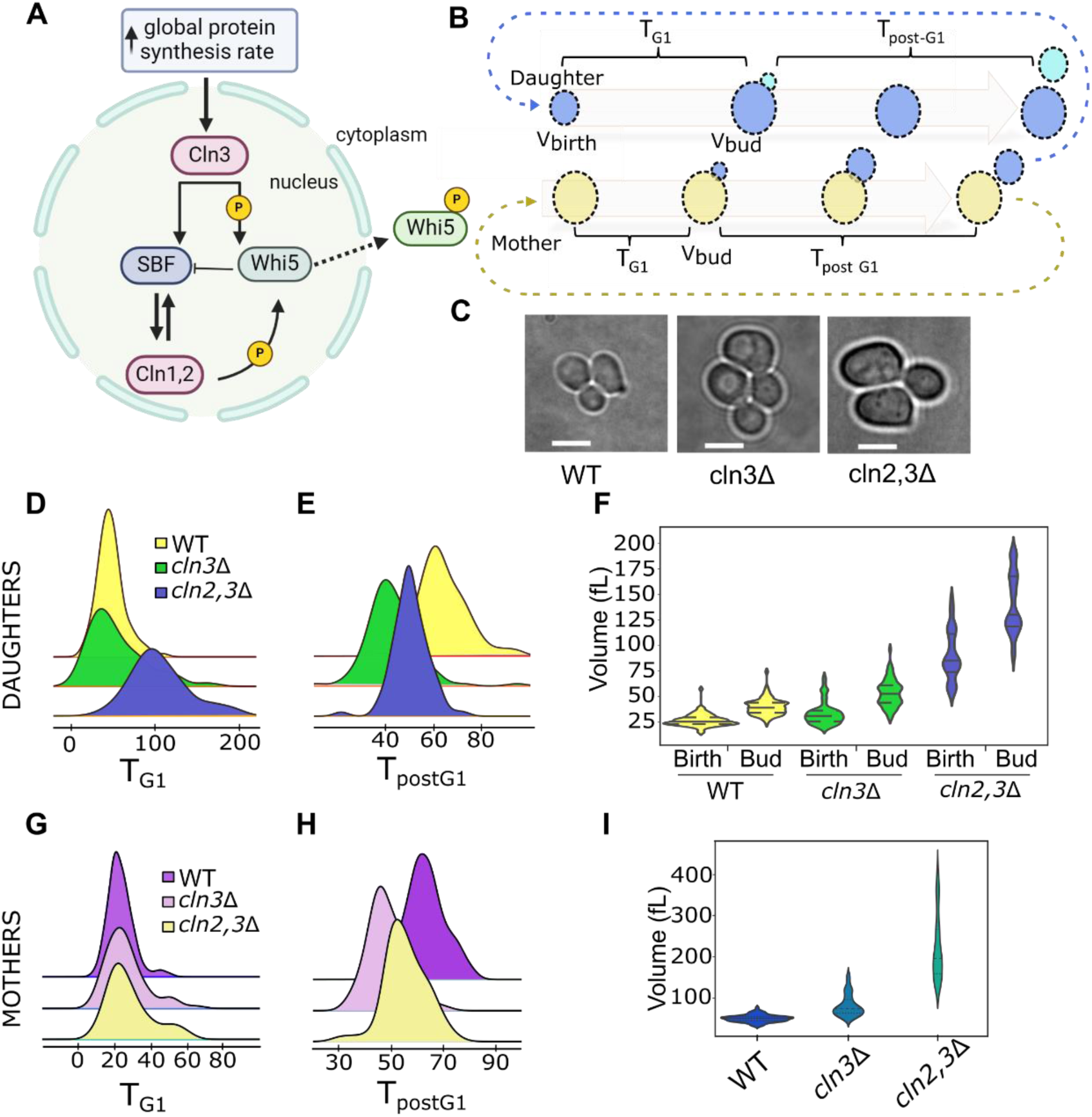
G1 cyclin mutants display differences in cell cycle progression and cell size. **A)** Schematic representation of the key players of the START transition network and their interactions. Cdk1 is the main CDK involved in cell cycle regulation in budding yeast (Enserink & Kolodner, 2010). During G1, Cdk1 activity is driven by three G1 cyclins, Cln1, Cln2 and Cln3 (functionally analogous to mammalian Cyclin D and E (Lew et al, 1991; Wang et al, 2009)). An increase in the expression of Cln3 (Litsios et al, 2019) leads to the formation of a Cln3/Cdk1 complex that eventually activates two major transcription factor complexes, SBF and MBF (Ferrezuelo et al, 2010; Wijnen et al, 2002). These complexes drive the transcription of Cln1 and Cln2, as well as hundreds of additional genes required for the G1/S transition (Bertoli et al, 2013; Teufel et al, 2019). Expression of Cln1 and Cln2 sets up a positive feedback loop in which Cln1/Cdk1 and Cln2/Cdk1 complexes phosphorylate and deactivate Whi5, a transcriptional repressor that binds and inhibits SBF. Deactivation of Whi5 triggers the G1/S transition and bud formation (Skotheim et al, 2008; Charvin et al, 2010). **B)** Illustration of the experimentally determined cell cycle phase durations and cell volumes for mother and daughter cells. **C)** Representative brightfield images of WT, *cln3Δ* and *cln2,3Δ* cells. Scale bar: 5μm **D)** Distributions of G1 durations (T_G1_) in WT (n = 109), *cln3Δ* (n = 97) and *cln2,3Δ* (n = 79) daughter cells. **E)** Distributions of post-G1 durations (T_postG1_) in WT (n = 109), *cln3Δ* (n = 97) and *cln2,3Δ* (n = 79) daughter cells. **F)** Volume distributions at birth (V_birth_) and budding (V_bud_) for WT (n = 82), *cln3Δ* (n = 77) and *cln2,3Δ* (n = 44) daughter cells. Median and 25th and 75th percentiles are displayed with dashed lines. **G)** Distributions of G1 durations (T_G1_) in WT (n = 97), *cln3Δ* (n = 71) and *cln2,3Δ* (n = 75) mother cells **H)** Distributions of post-G1 durations (T_postG1_) in WT (n = 97), *cln3Δ* (n = 71) and *cln2,3Δ* (n = 75) mother cells. **I)** Volume distributions at budding for WT (n = 70), *cln3Δ* (n = 76) and *cln2,3Δ* (n = 46) mother cells. Median and 25th and 75th percentiles are displayed with dashed lines.

Collectively, our results showed that, while loss of Cln3 does not lead to any pronounced changes in G1 duration in mothers or daughters, the loss of both Cln2 and Cln3 has a substantial impact on the G1 duration of daughters (Fig. 1D,G). Moreover, loss of G1 cyclins tends to cause a slight shortening of post-G1 phase durations in both mothers and daughters (Fig. 1E,H). Deletions of G1 cyclins result in cells that are larger than the WT as previously observed (Dirick *et al*, 1995; Teufel *et al*, 2019), with the double cyclin deletion resulting in the largest volume increase (Fig. 1 F,I). These observations will be used in the development of our optogenetic systems below.

### An optogenetic system for light-driven Cln2 expression: general design considerations

Our study of G1 cyclin deletion mutants showed that the transition from two functional cyclins to one causes a significant delay in daughter G1 progression. On the other hand, it is well-established that loss of the remaining cyclin leads to cell cycle arrest. It should therefore be possible to construct optogenetic systems that can either slow down or completely halt the cell cycle. To achieve these effects, one can conditionally express a G1 cyclin in a background where either one or two other G1 cyclins are deleted. Conditionally expressing any of the three G1 cyclins should be possible in theory, but we chose Cln2 because it has been found to exert stronger control on G1 progression than Cln1 (Queralt & Igual, 2004).

To regulate Cln2 transcription in a light-dependent manner, we used a previously developed optogenetic construct (Benzinger & Khammash, 2018) that consists of the light-sensitive EL222 transcription factor (Motta-Mena *et al*, 2014) fused to the VP16 transcriptional activation domain. Since the wild-type EL222 protein has a dark reversion time of 1-2 min (Motta-Mena *et al*, 2014) and Cln2 is an unstable protein (τ_1/2_ ∼ 10-15 min (Quilis & Igual, 2017)), sustained Cln2 expression over several minutes would require nearly continuous illumination, which would in turn limit the number of image fields that can be simultaneously stimulated and observed in microscopy experiments. We therefore used an EL222 mutant (EL222-AQTrip (Zoltowski *et al*, 2013)) which has a slower reversion time (∼30 min). The slow reversion of this mutant allows the use of brief (∼1 sec) light pulses every 5 min for establishing and maintaining sufficiently high Cln2 expression levels across a cell population. To maintain as uniform EL222 levels as possible, the VP16:EL222-AQTrip construct was integrated into the HO locus of our background strain, under the control of the constitutive ACT1 promoter.

Once activated by low-intensity blue (∼450-470 nm) light, EL222 homodimerizes and binds to its cognate binding site (Motta-Mena *et al*, 2014). Insertion of multiple EL222 binding sites to the promoter region upstream of a given target gene ensures sufficiently strong activation by EL222 (Benzinger & Khammash, 2018). To achieve optogenetic control of Cln2 expression, we replaced 700 bp upstream of the CLN2 ORF by an EL222-responsive promoter.

Cln2 is a weakly expressed protein (Dorsey *et al*, 2018), and it is well-established that Cln2 overexpression causes severe morphological and growth defects (McCusker *et al*, 2007; Lew & Reed, 1993; Barral *et al*, 1995). To avoid these artifacts while maintaining cell cycle control, an inducible Cln2 expression system would need to combine low maximal activity in the light (to keep Cln2 levels within physiological limits) with minimal leakage in the dark (to ensure Cln2 cannot promote cell cycle commitment without light stimulation).

To identify a promoter with the desired features, we considered a recently presented collection of EL222-responsive promoters for budding yeast (Benzinger & Khammash, 2018). Focusing on the two promoters at the low end of the expression scale, we chose pSPO13 and pGAL1 backbones for further testing. Spo13 is a meiotic regulator whose expression is tightly repressed during vegetative growth (Buckingham *et al*, 1990), while Gal1 expression is tightly repressed under glucose-rich conditions (Flick & Johnston, 1990). Despite this fact, cells harboring the pGAL1-based EL222 promoter showed signs of Cln2 overexpression (elongated shape and slow proliferation, Suppl. Fig. S2) even in darkness, suggesting that the leakage of this promoter is too high even under repressing conditions. On the other hand, we did not observe any signs of Cln2 overexpression in cells carrying the pSPO13-based promoter. We therefore chose this promoter for the construction of our optogenetic strains. Henceforth, we will refer to this light-dependent CLN2 construct as the OPTO-Cln2 system (Fig. 2A).

**Figure 2:**
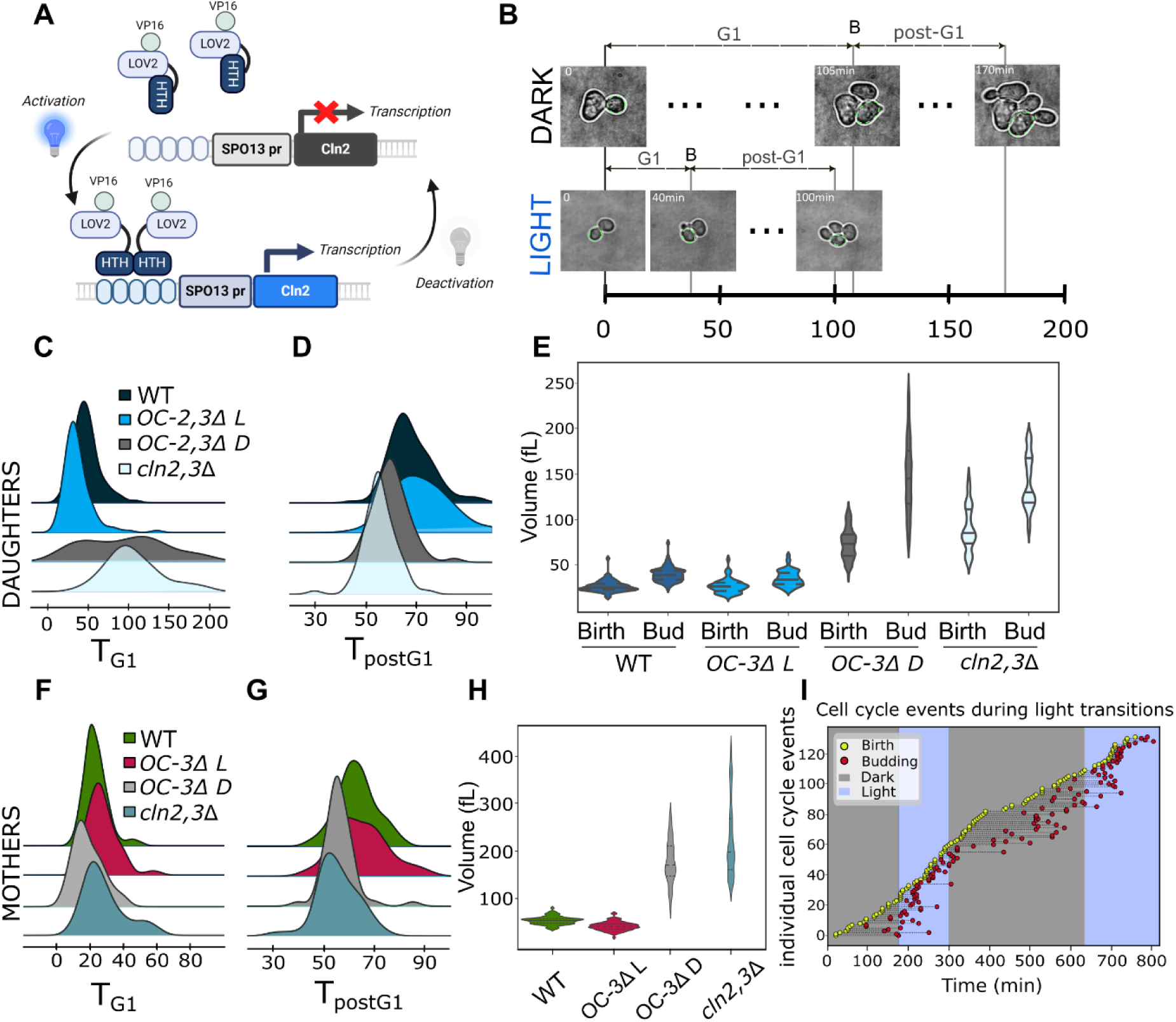
The OPTO-Cln2 system and dynamic regulation of G1 progression in an OPTO-Cln2 *cln3Δ* background. **A)** Activation of VP16-EL222 by low-intensity blue light triggers EL222 homodimerization and binding to its cognate binding sites. These sites are introduced upstream of the SPO13 promoter which replaces the endogenous CLN2 promoter. Exposure to blue light activates CLN2 transcription within a few minutes (Rullan *et al*, 2018). In the absence of light, EL222 spontaneously reverses to its dark state and is released from DNA, shutting down CLN2 transcription. The EL222 variant used here (AQTrip) has a dark reversion half-time of ∼30 min (Zoltowski *et al*, 2013). **B)** Phenotypic differences of OPTO-Cln2 daughter cells in light vs. dark conditions. Light activation leads to accelerated G1 progression and smaller cell volumes, while darkness increases both G1 duration and cell volumes. **C)** Distributions of G1 durations (T_G1_) in daughters of WT (n = 109), the OPTO-Cln2 *cln3Δ* (denoted OC-3Δ) strain in light (OC-3Δ L; n = 106) and darkness (OC-3Δ D; n = 61) and *cln2,3Δ* (n = 79). **D)** Distributions of post-G1 durations (T_postG1_) in WT (n = 109), OC-3Δ L (n = 106), OC-3Δ D (n = 61) and *cln2,3Δ* (n = 79) daughters. **E)** Volume distributions of daughter cells at birth (V_birth_) and budding (V_bud_) for WT (n = 82), OC-3Δ L (n = 63), OC-3Δ D (n = 104) and *cln2,3Δ* (n = 44). Median and 25th and 75th percentiles are displayed with dashed lines. **F)** Distributions of G1 durations (T_G1_) in WT (n = 97), OC-3Δ L (n = 75), OC-3Δ D (n = 51) and *cln2,3Δ* (n = 75) mother cells. **G)** Distributions of post-G1 durations (T_postG1_) in WT (n = 97), OC-3Δ L (n = 75), OC-3Δ D (n = 51) and *cln2,3Δ* (n = 75) mothers. **H)** Volume distributions in mother cells at budding for WT (n = 75), OC-3Δ L (n = 70), OC-3Δ D (n = 63) and *cln2,3Δ* (n = 46). Median and 25th and 75th percentiles are displayed with dashed lines. **I)** Cell cycle events of individual OC-3Δ daughters during light transitions. Cells were annotated for the moments of birth (yellow circles) and budding (red circles). In this way, the length of the dashed line segment between birth and budding corresponds to G1 duration. Grey and blue segments denote intervals of darkness and light exposure, respectively.

### The OPTO-Cln2 system enables dynamic control of G1 progression

Based on our characterization of G1 cyclin-deficient strains presented above, we first built a strain capable of transitioning between fast and slow cell cycle progression. To achieve this, we put Cln2 expression under light control in a strain lacking CLN3. Upon light exposure, we expected that this strain should divide as fast or faster than the WT due to continuous Cln2 synthesis which bypasses the requirement for Cln3 in early G1. Conversely, subjecting the cells to darkness should result in EL222 deactivation and Cln2 degradation, leaving Cln1 as the sole activator of the G1/S transition. This condition was expected to induce a phenotype resembling the *cln2Δcln3Δ* mutant.

Liquid cultures of OPTO-Cln2 *cln3Δ* cells had a population doubling time of 94 min under exponential growth in the dark, which is between the 89 min of the WT strain and the 100 min of the *cln2Δcln3Δ* mutant (Suppl. Fig. S1). On the other hand, light-exposed cultures of OPTO-Cln2 *cln3Δ* cells displayed a population doubling time of 71 min which is considerably shorter than the WT doubling time.

To determine in detail how the OPTO-Cln2 *cln3Δ* cells are affected by the presence or absence of light, we tracked the durations of G1 and non-G1 cell cycle phases of these cells with time-lapse microscopy, using WT and the *cln2Δcln3Δ* mutant as reference points. Dark-grown OPTO-Cln2 *cln3Δ* daughter cells had a prolonged G1 phase (100 min), similar to the *cln2Δcln3Δ* mutant and considerably longer than the average G1 duration of WT daughters (45 min) (Fig. 2C). Growth under low-intensity blue light caused a massive decrease in G1 phase duration of OPTO-Cln2 *cln3Δ* daughters (35 min), which now entered the cell cycle much sooner than WT daughters. On the other hand, the post-G1 duration of dark-grown OPTO-Cln2 *cln3Δ* daughters (60 min) was between that of the *cln2Δcln3Δ* mutant (55 min) and WT cells (65 min), and increased slightly (70 min) when these daughters were grown under blue light (Fig. 2D). Overall, the light-induced changes in daughter cell cycle progression indicated that the OPTO-Cln2 system in a *cln3*Δ background can strongly modulate G1 duration, producing daughters that divide as slow as *cln2*Δ*cln3*Δ or faster than WT.

Dark-grown OPTO-Cln2 *cln3Δ* mother cells did not display major changes between light and darkness, either in G1 (25 vs 15 min) or post-G1 duration (55 vs 65 min) (Fig. 2F, G). These phase durations were close to those of WT (G1: 20 min; post-G1: 65 min) and *cln2Δcln3Δ* cells (G1: 25 min; post-G1 55 min). Therefore, the OPTO-Cln2 system does not seem to affect cell cycle progression in OPTO-Cln2 *cln3Δ* mothers, as was already expected from the fact that WT and *cln2Δcln3Δ* mothers have similar cell cycle statistics.

As discussed above, besides affecting cell cycle progression, G1 cyclin deletions also affect the average sizes of both daughter and mother cells. To investigate whether the effect of the OPTO-Cln2 system induces cell volume changes in line with those expected from our results above, we quantified volumes of OPTO-Cln2 *cln3*Δ mothers at budding and daughter volumes at birth and budding, using the WT and *cln2Δcln3Δ* strains as reference points for light and dark conditions respectively. Light-grown OPTO-Cln2 *cln3Δ* daughters resembled WT daughters in terms of volume (birth: 26 fL vs 25 fL; budding: 34 fL vs 39 fL) (Fig. 2E). On the other hand, dark-grown OPTO-Cln2 *cln3Δ* daughters displayed a more than twofold increase in volume compared to WT (birth :73 fL; budding: 145 fL), in good agreement with the average volumes of *cln2Δcln3Δ* daughters (birth: 85 fL; budding: 130 fL). Light-grown OPTO-Cln2 *cln3Δ* mother cells were slightly smaller than WT at budding (40 fL vs 53 fL), while growth in darkness increased the average volume (170 fL) close to that of the *cln2Δcln3Δ* reference strain (201 fL) (Fig. 2H).

Taken together, our results indicate that the OPTO-Cln2 system in a *cln3Δ* background is capable of reproducing the *cln2Δcln3Δ* cell cycle and volume phenotypes in the dark. In the light, the system produces cells that are slightly smaller and divide faster than the WT, consistent with the notion that constitutive Cln2 expression fully compensates for the absence of Cln3, allowing cells to transition earlier into the S phase compared to WT cells. In summary, our tests demonstrated that the OPTO-Cln2 *cln3Δ* daughters can switch between a long and a short cell cycle, depending on the light condition, and that this effect is mostly attributed to the acceleration or slowdown of G1 phase (Fig. 2B). On the other hand, the cell cycle of mother cells remains largely unaltered in this optogenetic strain.

Following the characterization of the OPTO-Cln2 *cln3Δ* strain under steady-state growth, we tested how quickly cells respond to changes in light conditions. For this purpose, we used time-lapse microscopy to monitor the cell cycle progression of daughter cells, since this is the population that is affected by the light input. Our results revealed that daughters born shortly before or right after a darkness-to-light transition, exhibited a shorter G1 phase compared to daughters born in darkness (Fig. 2I). Conversely, daughters born shortly after a transition from light to darkness, displayed a longer G1 compared to daughters born in light. These findings suggest that Cln2 levels respond fast to light transitions, and this response drives changes in G1 duration for daughters that are born close to the transition times. In summary, the OPTO-Cln2 *cln3Δ* strain switches between fast- and slow-dividing daughter phenotypes, with fast transitions in either direction.

### The OPTO-Cln2 system generates light-dependent arrest and release

To develop a strain that can divide only in the presence of blue light, we introduced OPTO-cln2 in a *cln1Δcln3Δ* background. Contrary to the slow-dividing OPTO-Cln2 *cln3Δ* daughters, cell cycle progression of OPTO-Cln2 *cln1Δcln3Δ* is expected to be halted at the G1/S boundary when daughter and mother cells are shifted to darkness.

Liquid cultures of OPTO-Cln2 *cln1Δcln3Δ* cells grown under low intensity blue light had a population doubling time of 84 min, which was ∼10 min shorter than the doubling time of WT cells. This observation indicates that our optogenetic system can generate sufficiently high abundance of Cln2 to support fast proliferation even in a *cln1Δcln3Δ* background.

To determine if this strain undergoes arrest in the absence of light, a liquid culture growing in the presence of blue light was shifted to darkness while cell numbers were continuously monitored using flow cytometry. Following the shift to darkness, cell proliferation appeared to come to a halt within 1.5 to 2 hours, as was evident from the stabilization of cell counts (Suppl. Fig S3). This delay is expected since cells need to complete the cycle in which the light transition occurred and arrest in the G1 phase of the following cycle. This observation implies that Cln2 expression quickly drops below the threshold required to support proliferation when cells are moved to darkness.

More detailed characterization of this strain via time-lapse microscopy showed that the average G1 duration of daughter cells grown under blue light was nearly half of the average G1 duration of the WT cells (37 min vs 63 min), consistent with the fact that the presence of Cln2 bypasses the need for Cln3 upregulation in early G1 (Fig. 3A). Rapid passage through G1 was also evident in the volumes of these daughter cells, which budded at a smaller size compared to WT (27 fl vs. 38 fl) (Fig. 3C). A moderate but significant increase was observed in the post-G1 phase duration of these light-grown cells compared to WT (75 min vs 65 min) (Fig. 3B).

**Figure 3:**
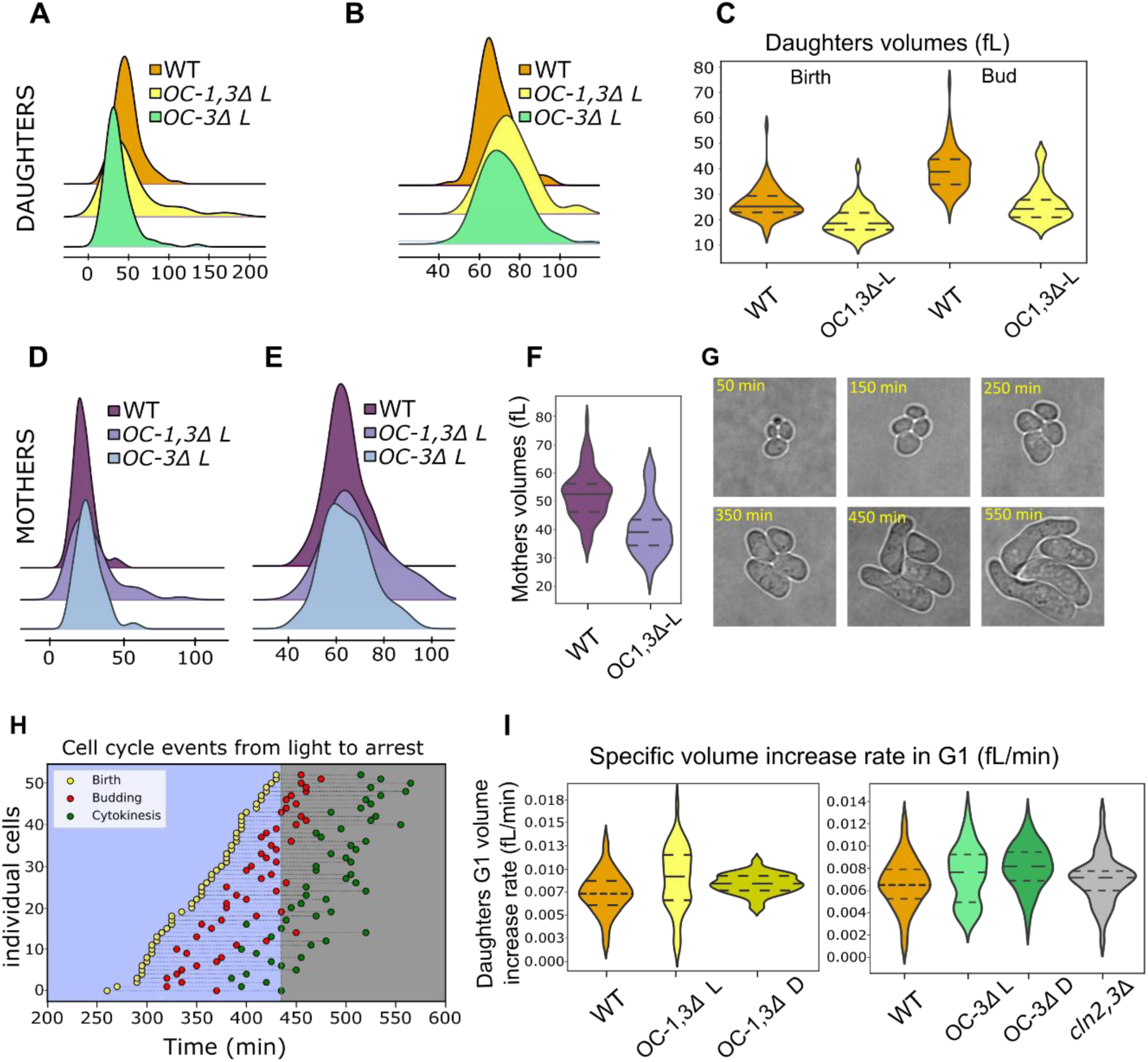
Dynamic regulation of cell cycle arrest and release with the OPTO-Cln2 system. **A)** Distributions of G1 durations (T_G1_) in WT (n = 109), OC-3Δ L (n = 72) and OPTO-Cln2 *cln1*Δ*cln3*Δ (denoted OC-1,3Δ) in light (OC-1,3Δ L; n = 106) daughters. **B)** Distributions of post-G1 durations (T_postG1_) in WT (n = 109), OC-3Δ L (n = 72) and OC-1,3Δ L (n = 106) daughters. **C)** Volume distributions of daughter cells at birth and budding for WT (n = 82) and OC-1,3Δ L (n = 48). Median and 25th and 75th percentiles are displayed with dashed lines. **D)** Distributions of G1 durations (T_G1_) in WT (n = 97), OC-3Δ L (n = 75) and OC-1,3Δ L (n = 57) mothers. **E)** Distributions of post-G1 durations (T_postG1_) in WT (n = 97), OC-3Δ L (n = 75) and OC-1,3Δ L (n = 57) mothers. **F)** Volume distributions of mother cells at budding for WT (n = 70) and OC-1,3Δ L (n = 38). Median and 25th and 75th percentiles are displayed with dashed lines. OC-1,3Δ D cells arrest unbudded in G1 **G)** Brightfield images of OC-1,3Δ light-grown cells, shifted to darkness at t=0 and grown under YNB + Glycose 2% agar pads for several hours. **H)** Cell cycle events of individual OC-1,3Δ daughters during light transitions. Cells were tracked for ∼8h during a transition from light to darkness and the cell cycle events (birth, budding and cytokinesis) of the last cycle before the arrest were annotated. Daughters that were born during the dark period and did not finish a cell cycle until the end of the experiment were excluded from the plot. **I)** Distributions of specific volume increase rates during G1 in daughters of WT (n = 82), OC-1,3Δ D (n = 64) and OC-1,3Δ L (n = 47) cells, OC-3Δ D (n = 103), OC-3Δ L (n = 57), and *cln2,3Δ* (n = 44). The WT distribution is presented twice to facilitate comparisons.

When grown under blue light, mother cells of the OPTO-Cln2 *cln1Δcln3Δ* strain displayed G1 and post-G1 phase durations that were similar to those of WT mothers (25 min vs 20 min for G1; 65 min for post-G1) (Fig. 3D, E). Moreover, the volumes of OPTO-mother cells were smaller compared to WT mothers (39 fL vs. 53 fL) (Fig. 3F). In brief, our results on light-grown OPTO-Cln2 *cln1Δcln3Δ* cells suggest that, in the presence of light, these cells proliferate similarly to light-grown OPTO-Cln2 *cln3Δ* and faster than WT cells, thanks to the shortened G1 duration of daughters.

When transferred to darkness, OPTO-Cln2 *cln1Δcln3Δ* cells arrest in the unbudded state (Fig. 3G). Moreover, as will be discussed below in detail, the transition from proliferation to arrest is fast (Fig. 3H). We observed that the arrested cells continued to grow in volume and acquired irregular shapes, while prolonged arrest (> 8 hours) led to cell death (Fig. 3G). This decoupling of volume dynamics from cell cycle activity prompted us to investigate more broadly whether the specific volume increase rate of OPTO-Cln2 daughters during G1 differs from the corresponding rate of WT and *cln2,3Δ* daughters. WT and the *cln2,3Δ* mutant displayed a small but non-statistically significant difference in their specific volume increase rates during G1 (0.0065 min^-1^ vs. 0.0072 min^-1^) (Fig. 3I). On the other hand, the specific volume increase rates of OPTO-Cln2 *cln1Δcln3Δ* daughters (in both light and darkness) displayed minor but statistically significant increases compared to WT daughters (dark-grown vs WT: 0.0071 min^-1^ vs. 0.0065 min^-1^; light-grown vs. WT: 0.0084 min^-1^ vs. 0.0065 min^-1^) (Suppl. Table 13). Similarly, a minor but significant increase was observed in dark-grown OPTO-Cln2 *cln3Δ* daughters (0.0082 min^-1^) compared to WT and *cln2,3Δ* (Fig.3I). In summary, OPTO-Cln2 *cln1Δcln3Δ* daughters grow slightly faster compared to WT during G1, while OPTO-Cln2 *cln3Δ* daughters show a small increase in growth rate only in darkness. Given the large differences in G1 progression among these strains, the observed differences in growth rates are quite small, suggesting that the mechanism(s) regulating volume changes during G1 operate largely independently of the cell cycle machinery.

### Differential regulation of G1 cyclins in OPTO-Cln2 strains alters TORC1/PKA activity dynamics

Proliferating cells need to accumulate biomass and increase in volume before dividing. In budding yeast, the cellular processes that lead to biomass increase are tightly regulated by highly conserved signaling pathways centered around the Target of Rapamycin Complex 1 (TORC1) (Loewith & Hall, 2011) and Protein Kinase A (Broach, 2012). These pathways respond to the availability, uptake and utilization of nutrients to balance anabolic processes that increase biomass with catabolic processes that turn it over (Broach, 2012).

To maintain homeostasis, cells need to couple the activities TORC1 and PKA with the cell cycle. As we showed recently (Guerra *et al*, 2022), this coupling results in fluctuations of the joint TORC1 and PKA activity in the cell cycle of WT mother cells. However, the extent to which the cell cycle machinery contributes to the generation of these fluctuations remains unclear. To assess this contribution, we used the OPTO-Cln2 system to alter G1 duration and observe how TORC1/PKA activity dynamics adapt to the change. We therefore monitored the joint TORC1 and PKA activity in our OPTO-Cln2 *cln3Δ* strains under light and dark conditions, focusing on daughter cells where the greatest changes in G1 duration are observed between light and dark conditions.

As a joint readout of TORC1 and PKA, we used the intracellular localization of Sfp1, a transcription factor that is targeted by both pathways (Guerra *et al*, 2022; Vuillemenot & Milias-Argeitis, 2022). Active TORC1 and PKA phosphorylate Sfp1 and promote its accumulation in the nucleus, whereas inactivation of either pathway leads to a relocation of Sfp1 to the cytoplasm (Lempiäinen *et al*, 2009; Jorgensen *et al*, 2004; Vuillemenot & Milias-Argeitis, 2022) (Fig. 4A). Monitoring the nuclear accumulation of GFP- and mNeonGreen-tagged Sfp1 during the cell cycle of individual WT mother cells had previously revealed two maxima, one in G1 and one in late G2, and we were able to associate the former with a peak in the joint TORC1 and PKA activity (Guerra *et al*, 2022). GFP is not appropriate for monitoring signaling activity in our optogenetic strains, since its excitation spectrum overlaps with the absorption spectrum of EL222. We therefore switched to a red fluorescent protein and monitored the localization dynamics of mScarlet-tagged Sfp1 using time-lapse microscopy (Fig. 4B). The nuclear-to-cytosolic ratio of Sfp1 was tracked during the cell cycle of both mother and daughter cells using a modification of our previously presented method to account for the lack of nuclear marker in our strains (Materials and Methods) (Suppl. Fig. S4).

**Figure 4:**
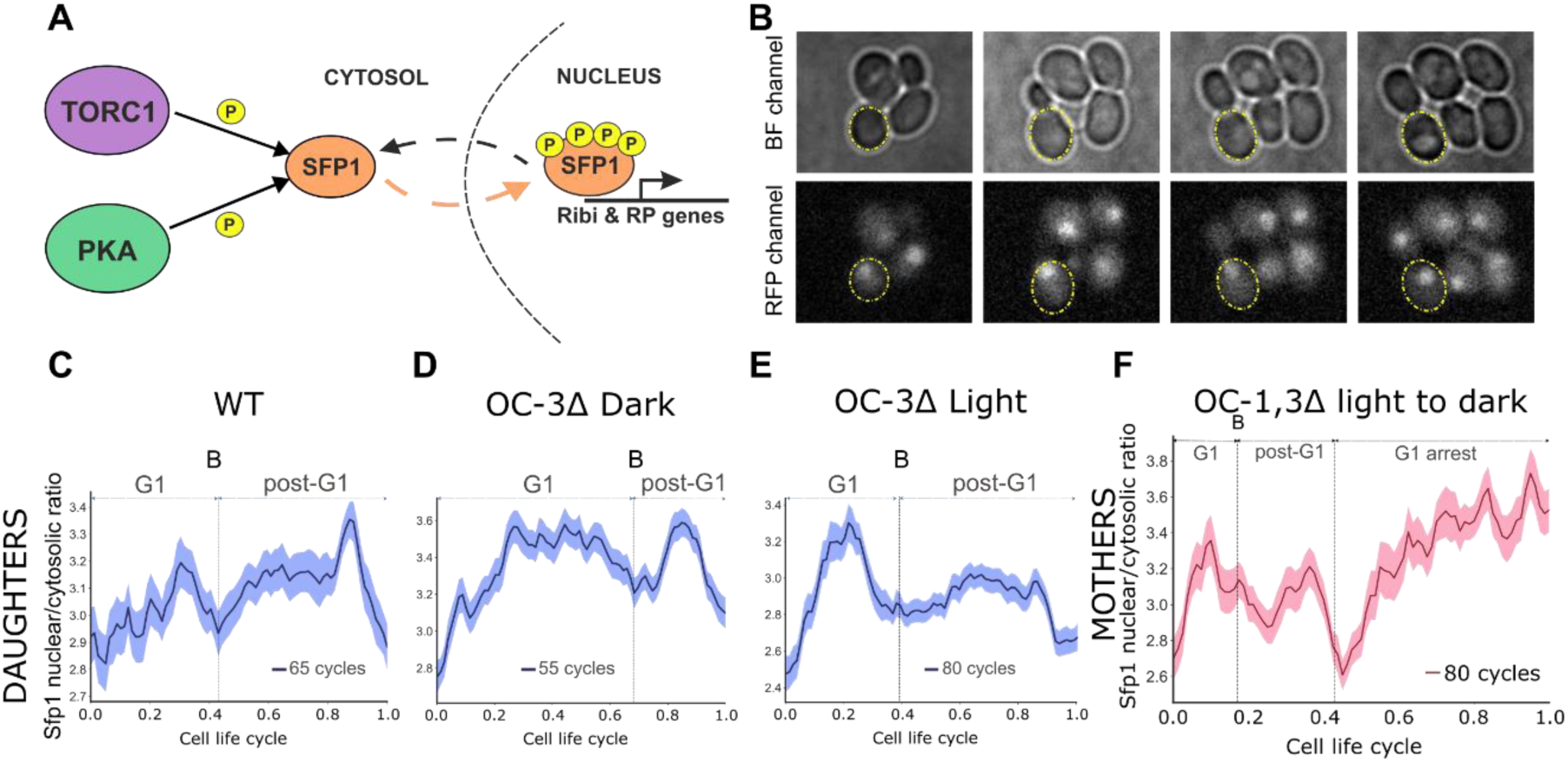
**A)** Schematic representation of the TORC1- and PKA-dependent regulation of Sfp1 phosphorylation and localization. **B)** Representative brightfield and fluorescent microscopy images used to track Sfp1-mScarletI localization during the cell cycle. Cells were segmented (yellow ellipsoids) and the ratio of nuclear to cytosolic Sfp1 was calculated as indicated in Methods. **C-E)** Nuclear-to-cytosolic ratio of Sfp1 in aligned cell cycles of daughter cells. Cell cycle events (birth, budding (B) and cytokinesis) were manually annotated in microscopy movies. The length of G1 and post-G1 phases in each plot was based on the ratio of average G1 and post-G1 durations to the average cell cycle duration of each strain. The bands denote the 95% confidence interval for the mean. **F)** Sfp1 nuclear to cytosolic ratio in light-grown OC 1,3Δ mother cells, following arrest in the dark. Cell cycles were aligned at four timepoints: cytokinesis, last budding(B), last cytokinesis and arrest. The length of each interval is proportional to the average duration of each phase relative to the total experiment duration.

As we had previously observed (Guerra *et al*, 2022), the localization of Sfp1-mScarlet displayed an oscillatory pattern during the cell cycle of WT mothers, with maxima at G1 and G2 and minima around budding and cytokinesis (Fig. 4G). This result confirmed that mScarlet can be reliably used to monitor Sfp1 localization during the cell cycle. Similar to WT mothers, mothers of the OPTO-Cln2 *cln3Δ* background displayed a G1 maximum under both under dark and light conditions (Suppl. Fig. S5).

Sfp1 did not exhibit a clear localization pattern during the G1 of WT daughters (Fig. 4C). Upon budding (i.e. entry to the cell cycle), the Sfp1 localization pattern of daughter cells became similar to that of mothers, with a maximum in G2 and a minimum around cytokinesis. Contrary to WT daughter cells, Sfp1 localization in dark-grown OPTO-Cln2 *cln3Δ* daughter cells displayed a substantial increase during G1 and a local minimum at budding (Fig. 4D). Light-grown OPTO-Cln2 *cln3Δ* daughters showed a G1 localization peak that was similarly delimited by cytokinesis and budding (Fig. 4E).

Collectively, these observations suggest that the combined TORC1 and PKA activity dynamics in OPTO-Cln2 *cln3Δ* daughters are mainly affected by changes in G1 progression. As G1 duration changes, the Sfp1 localization peak gets correspondingly adjusted, suggesting that the overall shape of TORC1 and PKA activity during G1 is mainly dictated by cell cycle-related processes.

We next asked how TORC1 and PKA respond when the cell cycle is halted. For this purpose, we tracked Sfp1 localization in OPTO-Cln2 *cln1Δcln3Δ* cells upon a transition from light to darkness. In this experiment, we followed mother cells that had budded shortly before or after the light was switched off, and aligned their traces at their cytokinesis, the last budding event, the following cytokinesis and the last time point at which each cell was tracked. Intriguingly, as cells transitioned from their last cytokinesis into G1 arrest, the nuclear-to-cytosolic (N/C) ratio of Sfp1 increased considerably, and was higher than the N/C ratio of the cell cycle preceding the arrest (Fig. 4F). On the one hand, this result indicates that in the absence of CDK activity, the combined activity of TORC1 and PKA remains high and even increases beyond its average during the cell cycle. On the other, it suggests that the observed minimum in TORC1 and PKA activity around budding in proliferating mother cells is most likely tied to S phase entry.

### Light-driven arrest induces synchronous release of budding yeast cells

Besides offering the possibility to study the (de)coupling of growth and division mechanisms, the uniform and rapid arrest of OPTO-Cln2 *cln1Δcln3Δ* cells can be used to generate synchronous cell populations, which can in turn enable the study of cell cycle-dependent processes in liquid cultures using different omics methods. Contrary to traditional methods of cell cycle synchronization in which only a single round of arrest and release is typically possible due to technical limitations (e.g. inducer washout), using light in principle allows multiple rounds of arrest and release, and thus better temporal resolution of cell cycle processes away from the point of release.

To establish the optimal conditions for light-induced synchronization experiments, we first determined the percentage of cells that arrest after a transition of a light-proliferating culture to darkness. For this test, we sampled liquid cultures at different time points during a transition from light to darkness, and calculated the percentage of budded vs unbudded cells via microscopic inspection. Unbudded cells accounted for the 89% of the population already after 120 min of darkness, which increased to 98% 180 min after the shift, demonstrating that our OPTO-Cln2 system can achieve a uniform cell cycle arrest (Fig. 5F).

**Figure 5:**
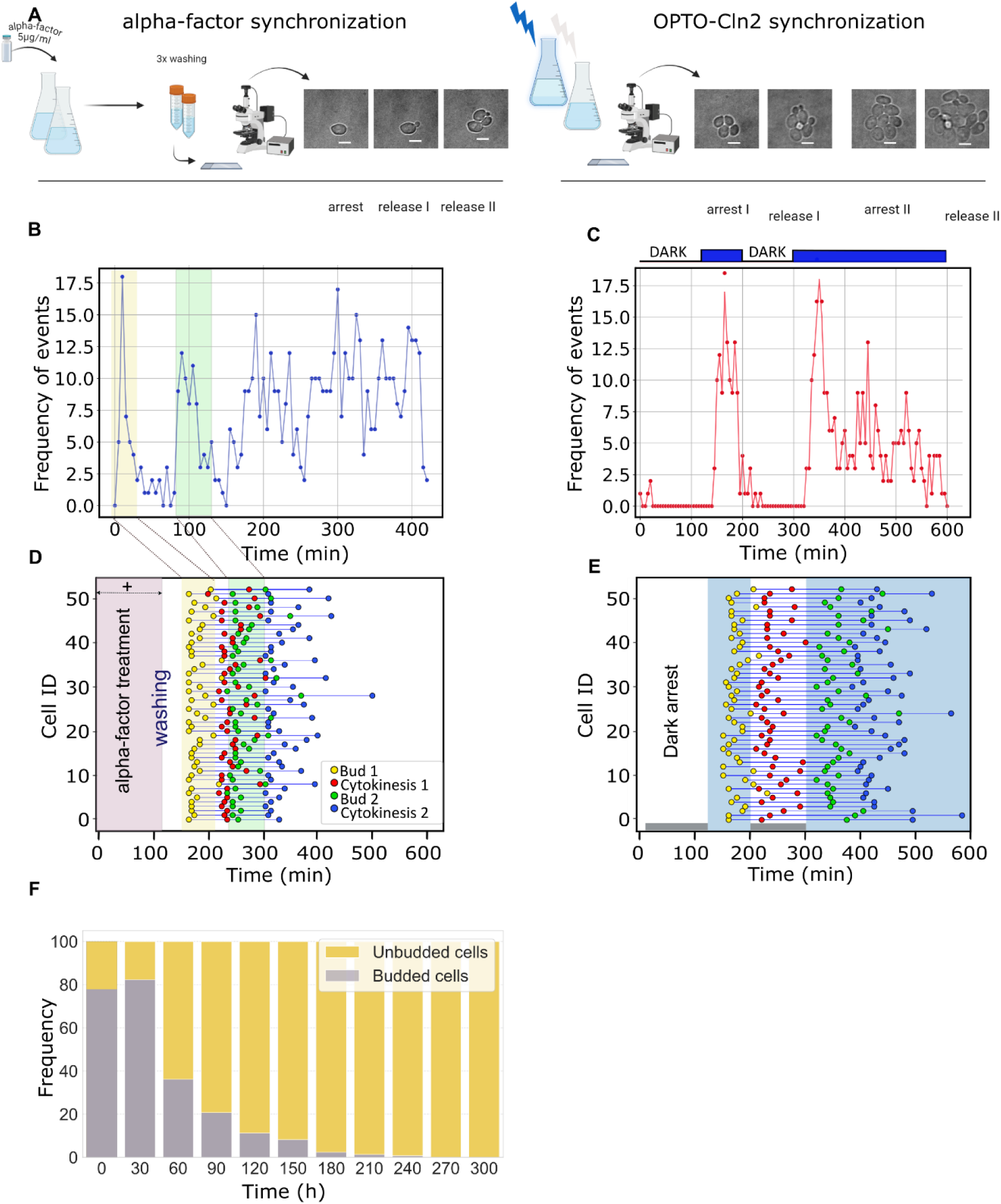
**A)** Schematic representation of experiments with alpha factor or light-induced synchronization. **B)** Frequency of budding events following alpha factor arrest and release plotted against time. **C)** Frequency of budding events following repeated light-induced arrest and release in OC-1,3Δ cells plotted against time. **D)** Cell cycle events of alpha-factor synchronized cells plotted against time, following two consecutive doublings after alpha-factor release. **E)** Cell cycle events of OC-1,3Δ cells during two consecutive light-induced arrests and releases plotted against time. **F)** Bar plot of percentages of budded (grey) versus unbudded (yellow) OC-1,3Δ fixed cells following arrest in the dark.

While longer dark periods increase the percentage of arrested cells, we noticed that prolonged arrest reduces the number of cells that can re-enter the cell cycle following a shift to light. Moreover, excessively long periods (>6 hours) of growth in darkness have adverse effects on cell morphology, as cells keep growing during the arrest and eventually die. To determine a dark period that optimizes the number of cells that can re-enter the cell cycle, we arrested cells grown in blue light under the microscope by shifting them to darkness for different lengths of time (80, 100, and 130 minutes) and calculated the percentage of budded cells upon release with another shift to blue light. We found that dark periods of 100 and 130 min both released ∼80% of the cells (Suppl. Fig. S7). On the other hand, we noticed that the 80 min arrest period was too close to the average cell cycle duration, which meant that a large fraction of cells had no time to reach G1 arrest before the light was switched on again. To reduce the effects of darkness on the cell morphology and enable multiple rounds of arrest and release, we proceeded with the optimized arrest period of 100 minutes.

To determine minimal period of the light induction that effectively releases the majority of cells for a single cycle before a second arrest, we performed experiments similar to the ones described above, with the periods of light and darkness reversed. These experiments established that 75 minutes of illumination are sufficient to release a large number of cells (∼ 90%) for a single cell cycle, while preventing entry to a second cell cycle (data not shown). Consequently, for repeated cycles of arrest and release, we used 100 minutes of darkness (arrest) followed by 75 minutes of light (release).

To benchmark the quality of synchronization achieved with our system, we compared its performance with the “gold-standard” arrest-and-release method for budding yeast, based on the alpha factor mating pheromone (Breeden, 1997; Amon, 2002). Addition of alpha factor to proliferating cells triggers a signaling cascade that ultimately leads to arrest in late G1 (Atay *et al*, 2016) (Fig. 5A). Subsequent washout of the alpha factor allows cells to re-enter the cell cycle in a highly synchronous manner. For our tests, we arrested WT YSBN6 cells by alpha-factor (5μg/ml) and OPTO-Cln2 *cln1Δcln3Δ* by a 100-min shift to darkness before their subsequent release (Fig. 5A). We then used time-lapse microscopy to assess the release at the single-cell level, by monitoring the timing of bud appearance in the released cells. For the alpha-factor released cells, buds started to appear ∼55 – 80 min after washout, with the majority of budding events occurring ∼60 min post-washout (Fig. 5B). For the OPTO-Cln2 *cln1Δcln3Δ* cells, we observed that buds started appear between 35 – 85 min following light induction, with the majority of budding events occurring ∼55 min post-induction (Fig. 5C). Overall, alpha factor and light induction displayed similar release dynamics.

To compare the degree of synchrony achieved by a single alpha factor release compared to two consecutive cycles of light-induced arrest and release, we determined the timing of cytokinesis and budding events of individual cells over time (Fig. 5D). While budding events appear within a narrow time window following alpha factor washout, cytokinesis events of the first post-release cycle become intertwined with budding events of the subsequent division cycle, making challenging the discrimination between the G2/M phase of the first cycle and the G1 phase of the subsequent cycle. On the other hand, repeated optogenetic arrest and release allows events from the one division cycle to remain distinct from those of the subsequent cycle (Fig. 5E).

Overall, our results demonstrate that the OPTO-Cln2 system can be used to induce population synchrony with quality comparable to that achieved by alpha factor, while avoiding the labor-intensive washout procedure required by the latter. Moreover, OPTO-Cln2 can induce repeat cycles of G1 arrest and release, providing better temporal resolution of late cell cycle events.

## Discussion

In this work, we employed a light-inducible system to dynamically perturb cell cycle progression in budding yeast through the regulation of the Cln2 G1 cyclin. To benchmark our system, we characterized the phenotype of single and double G1 cyclin deletion mutants, focusing on cell volume and the duration of G1 and post-G1 phases. Although evidence regarding these mutants is spread throughout literature, a systematic characterization of the double mutant was still lacking. Consistent with prior work (Tyers *et al*, 1993; Cross, 1988; Nash *et al*, 1988; Queralt & Igual, 2004; Teufel *et al*, 2019), our findings revealed an increase in cell size in the absence of one cyclin, which was even greater when two cyclins were deleted.

With regard to cell cycle progression, both the *CLN3* and *CLN2,3* deletions induced a slight shortening of post-G1 phases, while deletion of *CLN2,3* drastically increased the length of G1 in daughter cells. The observation that deletion of *CLN3* does not affect the timing of G1 progression in either mothers or daughters has been made before (Garmendia-Torres *et al*, 2018), and so has the shortening of post-G1 phases in this mutant. As we showed here, mothers of the *cln2Δcln3Δ* mutant display the same features. Besides providing reference points for the validation of our OPTO-Cln2 strains, these observations are interesting in their own right, as they cannot be explained by the established model of G1 regulation in budding yeast. Instead, they suggest the presence of compensatory mechanisms that alter the expression levels (or the timing) of the remaining G1 cyclins, possibly in connection with the increase in cell size. These mechanistic connections between cell size regulation and cell cycle commitment warrant further investigation.

Introduction of our OPTO-Cln2 construct in a *cln3Δ* or a *cln2Δcln3Δ* background allowed us to modulate the size and cell cycle progression of these strains in manner consistent with our expectations from the *cln3Δ* and *cln2Δcln3Δ* characterization results. Moreover, transitions between light-and darkness-associated phenotypes were fast and uniform across cell populations. In this respect, our OPTO-Cln2 system achieves a dynamic performance similar to that of a widely used methionine-inducible Cln2 system (Bean *et al*, 2006; Charvin *et al*, 2009, 2010; Perrino *et al*, 2021). An alternative, estradiol-inducible Cln1 system has recently been introduced (Ewald *et al*, 2016; Zhang *et al*, 2019; Irvali *et al*, 2023), but it has not been extensively characterized. In either case, the need to add or remove a nutrient or a hormone to alter cell cycle progression implies that these systems are better suited to microfluidics-based studies, where fast and repeated changes of the growth medium composition are possible. Moreover, removal and re-addition of a key nutrient such as methionine may also alter the activity of nutrient-sensing pathways such as TORC1. As we have demonstrated here, the OPTO-Cln2 system is compatible with both micro- and macroscopic (i.e. liquid culture) studies thanks to the use of light as an inducer. Moreover, the high sensitivity of the EL222 system to blue light (Benzinger & Khammash, 2018) allows the use of very low intensities (< 500 μW/cm^2^) that do not interfere with cell growth even over several hours. The AQTrip EL222 variant used here further reduces the minimal light requirements, as it can be activated by brief light pulses and remain active for several minutes.

The choice of promoter for driving Cln2 expression proved critical in our OPTO-Cln2 design, as maintaining low cyclin levels is essential for avoiding morphological artifacts (elongated, non-symmetric cells with an aberrant budding pattern) and slow proliferation. Cells appear to be quite sensitive to Cln2 levels, since even leaky expression from the pGAL1-based EL222 promoter in glucose was capable of inducing Cln2 overexpression in darkness. Similarly, deletion of the endogenous *CLN2* gene and introduction of a pSPO13-driven copy of *CLN2* in the X-1 integration locus (Mikkelsen *et al*, 2012) resulted in Cln2 overexpression (Suppl. Figure 6). On the other hand, a longer (∼1000 bp) truncation of the endogenous *CLN2* promoter was also found to be suboptimal, as it resulted in enlarged cells with highly variable daughter G1 duration (Suppl. Figure 7). These findings should provide valuable insights for the design of other inducible cyclin systems in the future.

An optogenetic cell cycle control system is an ideal tool for studying the coupling between the cell cycle machinery and the cellular processes that control growth. Our observation that the specific growth rate during G1 displayed only minor changes between WT, slow-dividing and even cell cycle-arrested daughters, suggests that processes leading to cell volume increase during G1 operate independently of the cell cycle machinery. On the other hand, our tests with Sfp1, a joint TORC1 and PKA readout, suggest that the activity of these pathways is coordinated with cell cycle events at the G1 boundaries: mother cells of both WT and optogenetic strains display a maximum of TORC1/PKA activity during G1 and minima at mitosis and budding, while OPTO-Cln2 *cln3Δ* daughters feature a similar localization pattern that scales with the length of G1 in dark vs. light conditions. Further evidence for the coupling of TORC1/PKA with the cell cycle is provided by arrested OPTO-Cln2 *cln1Δcln3Δ* cells, where TORC1/PKA activity remains high as cells keep increasing in size without entering the cell cycle.

Besides its utility in the study of coordination between growth and division, the OPTO- Cln2 system can be further used as an arrest and release device for the induction of population synchrony. Apart from alpha factor treatment, existing synchronization methods for budding yeast comprise drug treatments that activate cell cycle checkpoints, the use of temperature-sensitive mutants of essential cell cycle proteins, and centrifugal elutriation (Amon, 2002; Manukyan *et al*, 2011). Since drugs and temperature shifts can induce stress responses and alter cell physiology, alpha factor treatment and elutriation are considered the methods of choice for budding yeast. It should be noted that our background strain (the S88C-derived YSBN6) consistently displayed a lower degree of synchrony following alpha-factor release compared to CEN.PK (Suppl. Fig. S6), a strain commonly used in industrial applications, suggesting that alpha factor sensitivity is strain-dependent. More importantly, alpha factor triggers a major reorganization of gene expression and signaling via the mating response (Merlini *et al*, 2013; Liao & Thorner, 1980; Goranov *et al*, 2013). On the other hand, elutriation typically has a low yield of early G1 cells and requires special equipment (Rosebrock, 2017). Overall, both methods are technically challenging to apply and may provide variable results depending on user experience. Light-induced synchronization uses a non-toxic stimulus that specifically targets a single gene, and thus minimizes undesired side-effects of arrest-and-release. It is also easy to apply to large liquid cultures, which makes it possible to employ different omics methods to study large-scale changes in transcription, protein synthesis, phosphorylation and metabolism through the yeast cell cycle.

Given the good conservation of G1 progression mechanisms between yeast and mammalian cells (Morgan, 2007), we anticipate that the functional principle of our optogenetic system, which involves the transcriptional regulation of G1 cyclins, will be applicable to proliferating mammalian cells as well. More broadly, we expect that the ability to dynamically and reversibly modulate cell cycle progression will provide crucial insights into the mechanisms coordinating growth- and division-related processes.

## Materials and methods

### Plasmid and yeast strain construction

All yeast strains used in the study were constructed in the S288C-derived prototrophic genetic background YSBN6 ((Canelas *et al*, 2010), Euroscarf, Germany). CRISPR plasmids targeting the HO, *CLN1*, *CLN2* and *CLN3* loci were prepared based on a previously described protocol (Novarina *et al*, 2022). For the construction of each plasmid, we used pYTK-DN1 (containing the Cas9 cassette), pYTK-DN2 (spacer cassette), a double-stranded sgRNA and pYTK-DN5 (antibiotic resistance marker) from (Novarina *et al*, 2022). Individual sgRNAs were selected using the E-CRISPR tool following the guidelines of (Novarina *et al*, 2022) to ensure specificity. In the case of CLN2, the sgRNA was targeted at the promoter sequence to avoid modifying the coding sequence with synonymous mutations.

For the construction of the EL222 repair fragment (introduced into the HO locus), we modified plasmid pDB146 (Rullan *et al*, 2018) (kindly donated by Prof. Mustafa Khammash, ETH Zurich) by removing mScarlet-I, leaving ACT1pr-VPEL222_AQTrip-CYC1t. For the construction of the repair fragment containing an EL222-sensitive promoter upstream of the *CLN2* ORF, Gibson assembly was used to introduce EL222 binding sites (5x), the SPO13 core promoter (Benzinger & Khammash, 2018), a Kozak sequence (Benzinger & Khammash, 2018), the *CLN2* ORF and an mNeonGreen tag in a pFA6 backbone (mNeonGreen fluorescence of this construct was not clearly distinguishable from cellular autofluorescence and was therefore not used in this study). For the construction of Sfp1-mScarlet, Gibson assembly was used to create a linear fragment of pDB146 containing mScarletI and pFA6-SFP1-mNG-NATMX (Guerra *et al*, 2022) in order to substitute mNG part with mScarletI The plasmids containing the repair fragments were used as PCR templates, where long primers with tails homologous to the corresponding genomic integration locus were used to amplify the region of interest. All CRISPR target sequences and repair fragment tails used for genomic integration are listed in Supplementary Table 4. All genomically integrated constructs were verified by PCR and Sanger sequencing (Eurofins genomics, The Netherlands).

### Growth conditions and application of optogenetics

For all experiments, yeast cells were grown in YNB medium (Formedium) supplemented with amino acids and 2% glucose. All liquid cultures were cultivated at 30°C with shaking at 300 rpm. To ensure exponential growth at the start of our experiments, precultures were appropriately diluted in the evening prior to each experiment, so that cells would be in mid-log phase (OD <1) on the day of the experiment. For experiments starting from dark conditions, precultures were grown in flasks covered with aluminum foil, and cell sampling was conducted in a dark room illuminated with a red LED light. For light-grown precultures, illumination was provided by a breadboard carrying 3 or 4 blue LEDs (Lumex, part no. SSL-LX5093USBC, 470 nm) that was attached to the incubator wall facing the preculture flask (Supplementary Figure 6).

### Microscopy

Microscopy experiments were performed using inverted fluorescence microscopes (Eclipse Ti-E, Nikon Instruments). Temperature was kept at 30 °C with the help of a temperature-controlled microscope incubator (Life Imaging Services). All images were obtained with a 100x Nikon S Fluor (N.A. = 1.30) objective and were recorded using iXon Ultra 897 DU-897-U-CD0-#EX cameras (Andor Technology). For fluorescent microscopy, LED-based excitation systems (pE2; CoolLED Limited and AURA; Lumencor) were used. For red fluorescent protein (mScarlet-I) measurements, cells were excited at 565 nm (excitation filter: 540–580 nm, dichroic: 590 nm, and emission filter: 600–650 nm). To activate the EL222 system through the microscope, we used a LED light source (CoolLED pE2) centered at 440 nm at 1% intensity, illuminating the cells for 1 sec every 5 min. During brightfield imaging, a long-pass (600 nm) filter was used to prevent unwanted activation of the EL222 system. The Nikon perfect focus system was used to prevent loss of focus in all experiments, and image acquisition was performed every 5 min.

For imaging, mid-log yeast cells grown on YNB supplemented with amino acids and 2% glucose were spotted on a microscopy slide and covered with an agarose pad (YNB +aa +2% glycose + 1% agarose). Whenever necessary, sample preparation for microscopy was performed in a dark room using a red-light LED lamp to avoid sample exposure to ambient blue light. Prior to each experiment, the samples were kept at the microscope for ∼1−2 h for cells to adapt to the new growth environment. For experiments requiring light-grown precultures, cells that were transferred to the microscope were kept illuminated by an LED breadboard (described above) during the adaptation period (Supplementary Figure 6).

### Image Analysis

All brightfield images were contrast-enhanced to facilitate segmentation, and fluorescent images were background corrected using the rolling ball subtraction plugin of ImageJ. For cell segmentation and tracking in time-lapse experiments, brightfield images were processed with the semi-automatic ImageJ plugin BudJ (Ferrezuelo *et al*, 2012). BudJ was used to provide estimates of cell volumes, as well as segmentation masks for processing the fluorescent channel images. Analysis of the BudJ-generated data and fluorescent images was performed using a custom-made script Python script.

Due to the fact that the absorption spectrum of EL222 extends up to 500 nm (Nash *et al*, 2011) and Sfp1 was tagged with a red fluorescent protein (mScarlet-I), we could not use a nuclear marker as in our previous work (Guerra *et al*, 2022) to accurately locate the yeast nucleus. We therefore used a modified procedure for the estimation of nuclear and cytoplasmic fluorescence, in which the fluorescence intensity of Sfp1 itself (which is always partially nuclear in rich nutrients) was used to define an area of brightest pixels via local thresholding (threshold_local() from the skimage library) within each cell mask.

By varying the offset parameter of the local threshold for each cell, we determined a bright region that contains no more than 30% of the total cell pixels, a percentage that is slightly larger than the average fraction of nuclear to whole-cell area (Jorgensen *et al*, 2007). The resulting region was then eroded using a 3x3 square structuring element to avoid inclusion of small non-connected regions of bright pixels, leaving only a region of contiguous pixels that should be located within the nucleus (nuclear ROI). The nuclear intensity of Sfp1 was then estimated from the average fluorescence of a circular region at the center of the nuclear ROI with a radius of three pixels. To estimate the cytosolic signal, a larger concentric circle with a radius of nine pixels was defined to ensure exclusion of nuclear pixels, and the average fluorescence of pixels between the cell boundary and the boundary of this larger circle was calculated. The average nuclear and cytosolic fluorescence were in turn used to calculate the nuclear-to-cytosolic ratio of Sfp1.

### Determination of cell cycle phase durations

To estimate the doubling times and (post-)G1 durations of individual cells, a custom-made ImageJ plugin (Click_cells2) was used to manually annotate events (birth, budding, cytokinesis and death) for tens of cells throughout a microscope movie. The output of the ImageJ plugin was further analyzed using a Python script to calculate statistics and make plots. The annotation of individual events was based on visual inspection of cellular features. Birth of daughter cells (and cytokinesis) was assigned to the time point at which a dark segment was visible between the mother and daughter cell, implying that cytokinesis had been completed. Budding was assigned to the time-point at which a dark spot, indicating the nascent bud, was visible on the periphery of a cell. The G1 duration of a daughter cell was then defined as the length of the interval between its birth and first budding event. Likewise, the G1 duration of a mother cell was defined as the length of the interval between a cytokinesis event and the following budding event. The doubling time of a mother cell was calculated as the length of the interval between two consecutive budding events. Finally, the S/G2/M (or post-G1) duration of a cell was defined as the length of the interval between budding and the following cytokinesis event. Note that, due to the use of a 5-min sampling interval (cf. Microscopy section), the medians of all durations reported in the main text are multiples of 5.

### Measurement of the Sfp1–mScarlet nuclear-to-cytosolic ratio in single cells

To compare and average the Sfp1 N/C ratio during cell cycles of different lengths, individual cell cycle traces were aligned at three cell cycle events: cytokinesis (birth for daughters), budding (end of G1), and the subsequent cytokinesis event. The traces were interpolated with the same number of time points on an axis representing cell cycle progression, by allocating to each phase (G1 and post-G1) a number of points proportional to the average duration of that phase in a given population. The aligned and interpolated traces were subsequently averaged to obtain the localization patterns presented here

### Light-induced synchronization (with cell fixation)

Log-phase (OD< 1) light-grown cultures were covered with aluminum foil to induce cell cycle arrest and were kept in the 30 C incubator. Sample collection was performed every 30 min, by pelleting 1 ml of culture, resuspending in ice-cold 4% formaldehyde fixation solution, and washing in ice-cold PBS. Individual time point samples were stored at 4 °C prior to microscopic quantification. For this quantification, 2 μl of each sample were spread on a microscopy slide and covered with an agar pad infused with 1x PBS solution and 1% agarose. Acquisition of multiple fields per sample was performed using the custom multipoint definition in NIS Elements, set to scan an area of 10x10 fields using 1% overlap, which resulted in 100 images for each sample. The percentage of budded vs. unbudded cells in each image was then calculated by visual inspection.

### Alpha-factor treatment

Before the addition of alpha factor, precultures of exponentially growing WT cells were diluted to OD 0.2 in 20ml YNB supplemented with amino acids and 2% glucose. A stock of 0.5 mg/ml alpha-factor (Eurogentec, Belgium) was diluted to a working concentration of 5 μg/ml before the addition in the culture. After alpha factor addition, the cultures were left at 30 °C for 2h. Subsequently, the alpha factor was washed to release the cells into the cell cycle. For alpha factor wash-off, cells were centrifuged (2000 rpm for 2 min) and resuspended 3 times in 2x volume of YNB medium without alpha-factor before being resuspended in 1x volume of YNB. For fixation experiments, samples were collected every 30 min as described above. For the microscopy experiments with alpha factor treatment, cells were kept in the incubator without disturbance for the duration of the treatment (2h). Following alpha factor washout (time zero in Fig. 5C), microscopy experiments were performed in YNB-agar pads as indicated above.

## Acknowledgements

We would like to thank Prof. Mustafa Khammash (ETH Zurich, Switzerland) for providing the pDB146 plasmid and Michiel Punter (University of Groningen) for the cell cycle annotation plugin for ImageJ.

A.K. and A.M.-A. were supported by the Dutch Research Council (NWO) through an NWO-VIDI grant to A.M.-A. (project number 016.Vidi.189.116).

