## Supplementary Figures and Tables for "Rapid and reversible regulation of cell cycle progression in budding yeast using optogenetics"

A

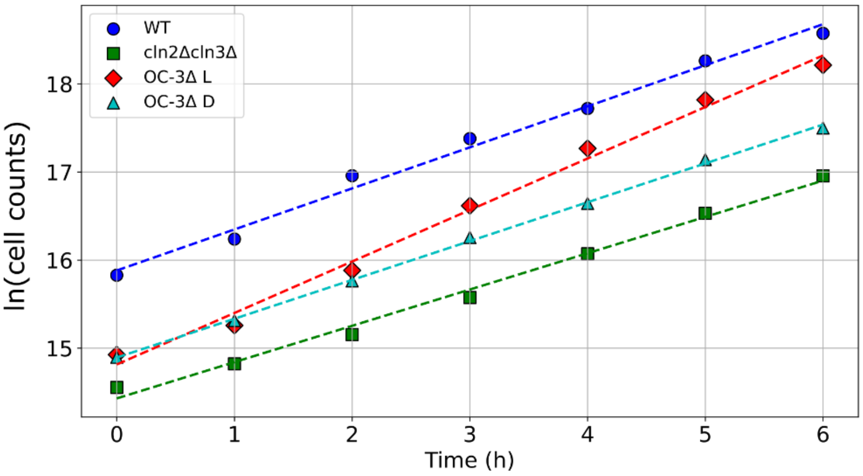


| Strains | k | T | T(min) |
|---|---|---|---|
| WT | 0.466 | 1.4874403 | 89.2464181 |
| OPTO-cln2_Light | 0.584 | 1.18689586 | 71.21375143 |
| OPTO-cln2_Dark | 0.4409 | 1.5721188 | 94.32712822 |
| cln2_3Δ | 0.4122 | 1.68157977 | 100.8947861 |

B C

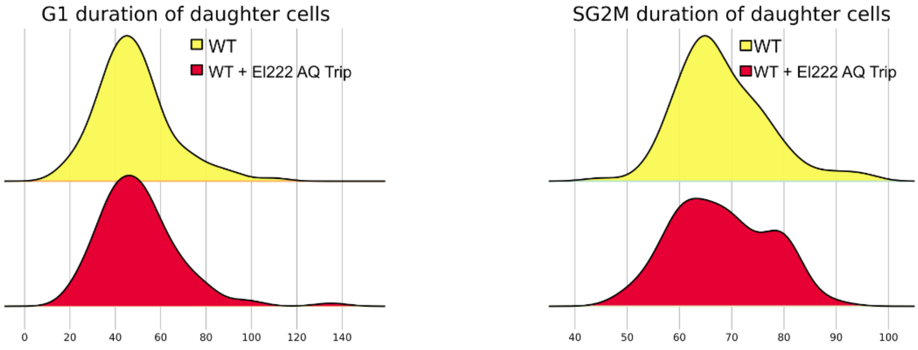


**Supplementary Figure 1**. A) Growth curves of WT, *cln2Δcln3Δ* and OPTO-Cln2 strains grown in YNB medium supplemented with amino acids and 2% glucose. OPTO-Cln2 liquid pre-cultures were grown in light or dark conditions prior to each experiment. The table displays the growth rate (*k*) and the population doubling time (calculated from ln(2)/k) of every strain. B) Distributions of G1 durations of WT daughters and of daughters expressing the VP-EL222 construct on the HO locus (without the EL222-responsive promoter). C) Distributions of post-G1 durations of WT daughters and of daughters expressing the VP-EL222 construct on the HO locus (without the EL222-responsive promoter).

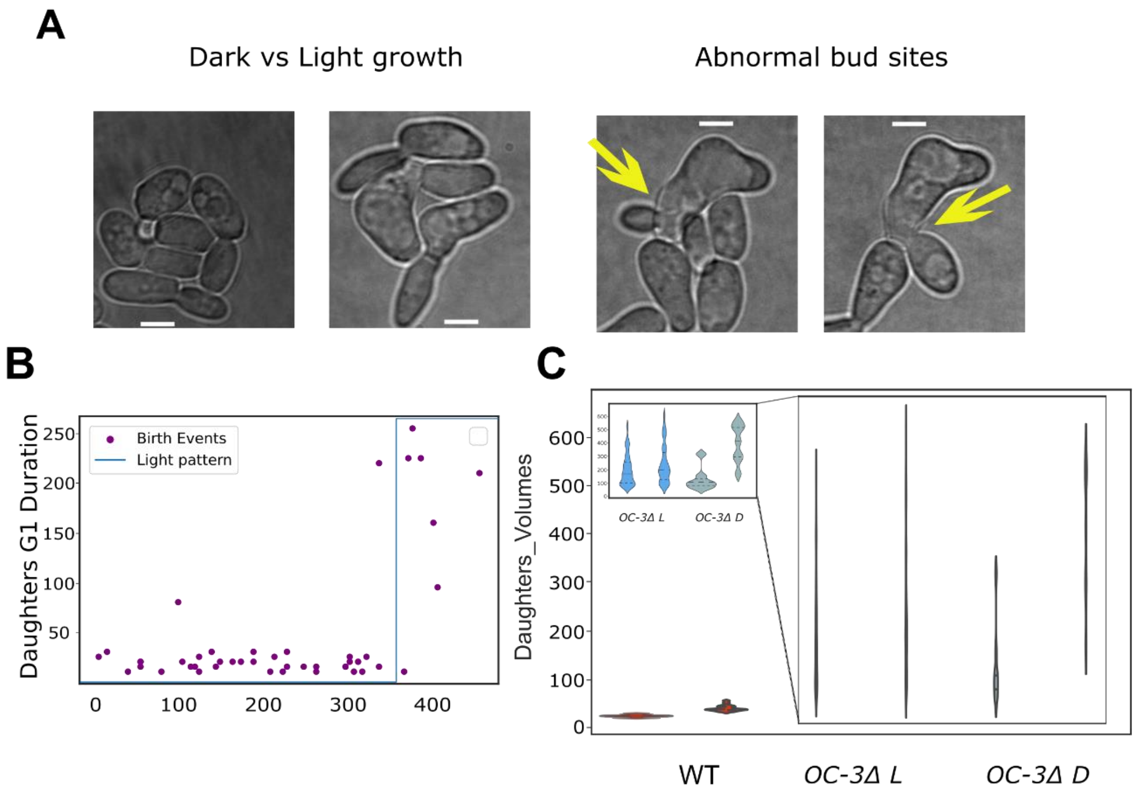


**Supplementary Figure 1. pGAL1 OPTO-Cln2 cells display phenotypic defects (large size and abnormal budding patterns) due to Cln2 overexpression.** A) Representative brightfield images of pGAL1 OPTO-Cln2 cells. Left image pair: comparison of cells grown in darkness vs light Right image pair: examples of abnormal budding patterns under light (right). B) Scatter plot of G1 durations in pGAL1 OPTO-Cln2 daughter cells during a transition from darkness to blue light. The x-axis denotes the birth moment of each tracked daughter, while the y-axis displays the corresponding G1 duration. Contrary to the expectation, daughter G1 durations are relatively short in darkness and increase massively when cells are shifted to light. C) Volume distributions of daughter cells at birth (left plot) and budding (right plot) for WT (n = 17), pGAL1 OC-3Δ L (n = 43), pGAL1 OC-3Δ D (n = 45). Median and 25th and 75th percentiles are displayed with dashed lines.

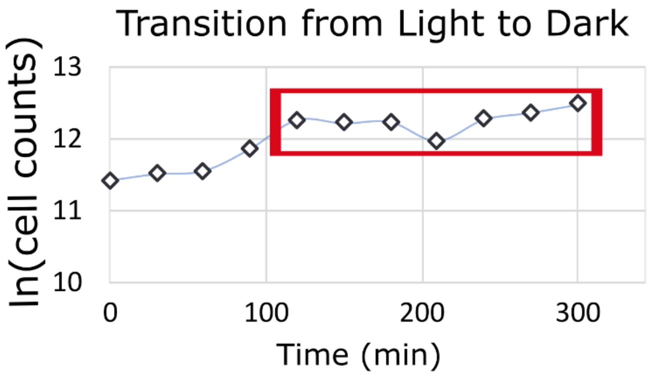


**Supplementary Figure 2**. Growth curve of OC-1,3Δ cells in YPD transitioning from a light-grown preculture that was shifted to darkness at t = 0. Post-G1 cells that encounter the light transition at t = 0 need to finish their ongoing cell cycle, which results in an increase in cell numbers ~100 min after the switch. From 100 min onwards, the cell counts stabilize, indicating arrest of cell cycle progression.

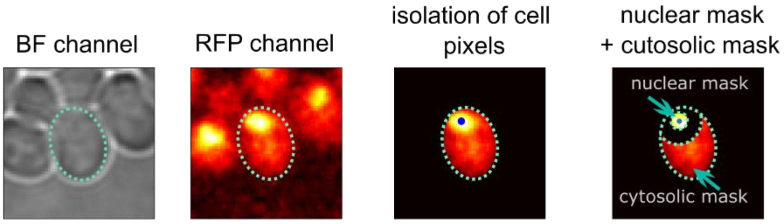


**Supplementary Figure 3**. Schematic representation of the pipeline used to analyze the N/C ratio of Sfp1-mScarlet-I. The cell segmentation produced with the BudJ plugin in the brightfield was used to define a cell mask in the RFP channel. The pixles of the mask were processed with the help of local tresholding (Materials and Methods) to identify the brightest region in the image whose area does not exceed 30% of the total cell area (nuclear ROI). Nuclear Sfp1 intensity was calculated from a smaller region within the nuclear ROI. A larger mask with the same center (black area) was defined to extend beyond the nucleus, and the remaining cell pixels were used to defined a cytosolic ROI. The Sfp1 N/C ratio was calculated by dividing the average intensity of the nuclear ROI by the average intensity of the cytosolic ROI.

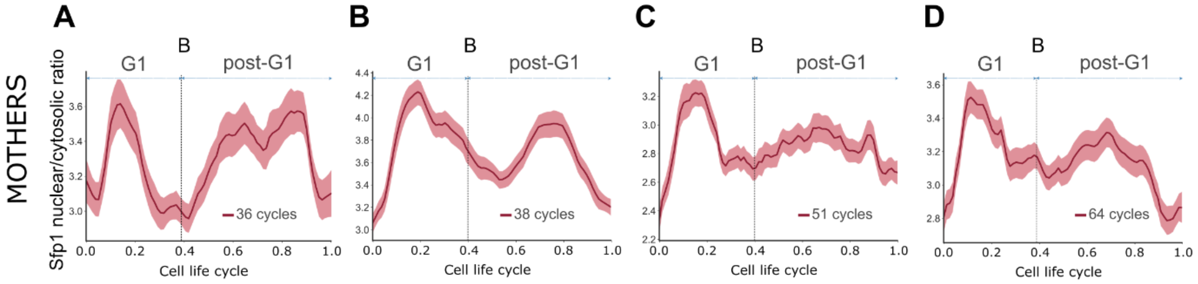


**Supplementary Figure 5**: **A-D)** Nuclear-to-cytosolic ratio of Sfp1 in aligned cell cycles of mother cells. Cell cycle events (cytokinesis, budding (B) and the following cytokinesis) were manually annotated in microscopy movies. The length of G1 and post-G1 phases in each plot was based on the ratio of average G1 and post-G1 durations to the average cell cycle duration of each strain. The bands denote the 95% confidence interval for the mean. Consistent with our observation that light does not alter cell G1 duration in OPTO-Cln2 cln3Δ mother cells, no major changes were observed in the G1 localization profile of Sfp1 in these cells. On the other hand, the attenuated localization peak of Sfp1 in G2 warrants further investigation.

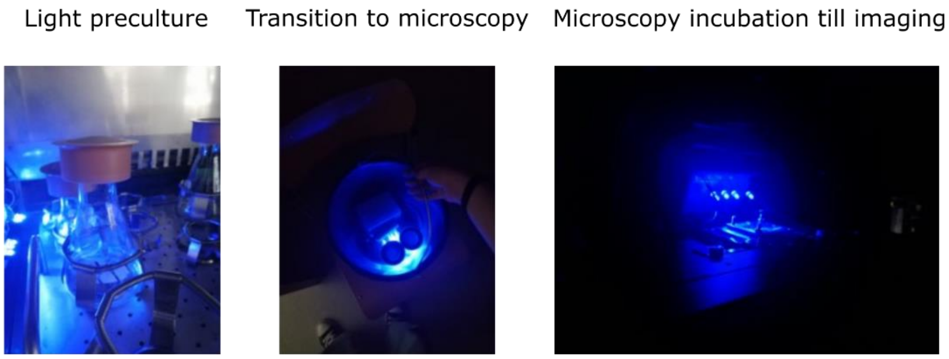


**Supplementary Figure 6:** Images showing the setup of the light illumination of liquid cultures of OC-1,3Δ cells in the rotating incubator (left), their transition to the microscopy room (middle) and during the waiting time (1-2h) in the microscopy chamber, before the beginning of the experiment.

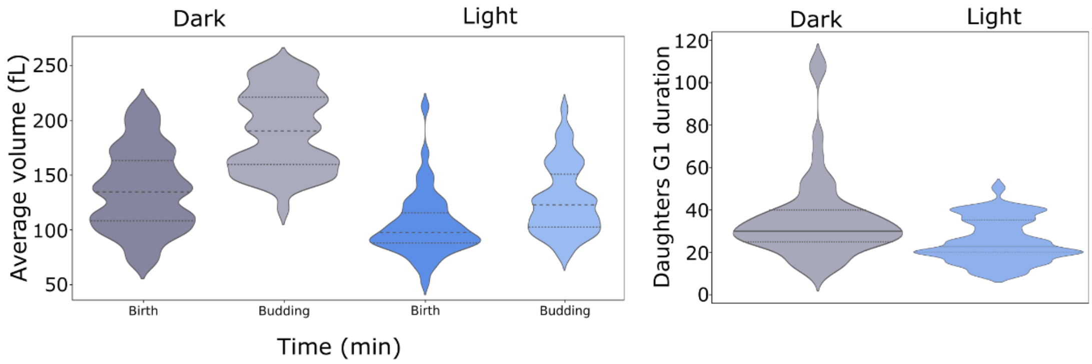


**Supplementary Figure 7:** The OPTO-Cln2 construct integrated in the X1 locus results in overexpression of Cln2, as suggested by the large volume of daughter cells during birth and budding in both light conditions (left) and the very short G1 duration in both darkness and light (right).

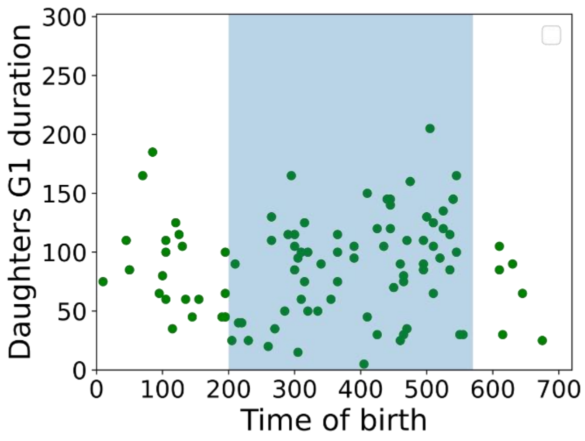


**Supplementary Figure 8:** The OPTO-Cln2 construct featuring a larger (1000 bp) truncation of the endogenous CLN2 promoter resulted in large heterogeneity in the G1 duration of daughters s in both dark and light conditions. Scatter plot of individual daughter G1 durations plotted at the moment of daughter birth during transitions between darkness (white) and light (blue) conditions.

**Supplementary Table 1:** List of strains used in the study.

| | | |
|---|---|---|
| YSBN6 | *MAT**a** ho::HphMX4* | (Canelas et al., 2010) |
| yAK29.2.3α | YSBN6 *ho::prACT1-VP-EL222-CYC1-hphMX4* | This study |
| yAK31.1.1α | YSBN6 *ho::prACT1-VP-EL222-CYC1-hphMX4, cln2Δ, cln3Δ* | This study |
| yAK38.1.6α | YSBN6 *WHI5-mCherry:Ble, ho::prACT1-VP-EL222-CYC1-hphMX4, El222(5xBS)-prSPO13-CLN2-mNG, cln3Δ* | This study |
| yAK35.1.3α | YSBN6 *Whi5:WHI5-mCherrry:Ble, ho::prACT1-VP-EL222-tCYC1-hphMX4, cln3:cln3Δ* | This study |
| yAK61.1.5α | YSBN6 *WHI5-mCherry:Ble, ho::prACT1-VP-EL222-CYC1-hphMX4, El222(5xBS)-prSPO13-CLN2-mNG, cln3Δ, cln1Δ* | This study |

| | | |
|---|---|---|
| yAK71.2.3γ | YSBN6 *WHI5-mCherry:Ble, ho::prACT1-VP-EL222-CYC1-hphMX4, El222(5xBS)-prGAL1-CLN2-mNG, cln3Δ* | This study |
| yAK76.1.4 | YSBN6 *ho::HphMX4, SFP1-mScarletI-NatMX* | This study |
| yAK76.2.4 | YSBN6 *ho::prACT1-VP-EL222-CYC1-hphMX4, El222(5xBS)-prGAL1-CLN2-mNG, cln3Δ, SFP1-mScarletI-NatMX* | This study |
| yAK76.3.4 | YSBN6 *ho::prACT1-VP-EL222-CYC1-hphMX4, El222(5xBS)-prGAL1-CLN2-mNG, cln3Δ, cln1Δ, SFP1-mScarletI-NatMX* | This study |

**Supplementary Table 2:** List of plasmids used in the study.

| Plasmid | Insert | Integration | Source |
|---|---|---|---|
| | EL222(5xBS)-prSPO13-Kozak-CLN2-mNG | CLN2 locus | This study |
| pSC1.1.3 | GAL1(UAS)-EL222(5xBS)-prGAL1-CLN2-mNG | CLN2 locus | This study |
| pLV1 | NLS-VP16-EL222 AQTrip | HO locus | |
| pYTK123 | pPGK1 - Cas9 - tPGK1 + HO-sgRNA1 | - | This study |
| pYTAK140 | pPGK1 - Cas9 - tPGK1 + CLN2-sgRNA1+CLN3-sgRNA2 | - | This study |
| pDB146 | ACT1pr-mScarletI-NLS-VP16-EL222_AQTrip-CYC1t | | Rullan *et al*, 2018 |
| Sfp1_mNG_Nat | pFA6-Sfp1_CDS-mNeonGreen-NatMX-Sfp1_DOWN | SFP1 locus | Guerra *et al*, 2021 |
| Sfp1-mScarlet | pFA6-Sfp1_CDS-mScarletI-NATMX-Sfp1_DOWN | SFP1 locus (SFP1 mScarletI) | This study |

**Supplementary Table 3:** List of primers used in the study.

| **Primer** | **Sequence** | **Application** |
|---|---|---|
| EL222-prSPO13_fwd | CGGCATCAGAGCAGATTGTAGAAG | Construction of GA pFA6-El222(5xBS)-prSPO13-cln2-mNG |
| EL222-prSPO13_rev | cactagccatAATTATTCTCGACTCAACTTCAATCCG | Construction of GA pFA6-El222(5xBS)-prSPO13-cln2-mNG |
| GA-cln2_fwd | gagaataattATGGCTAGTGCTGAACCAAG | Construction of GA pFA6-El222(5xBS)-prSPO13-cln2-mNG |
| GA-cln2_rev | ccatagaaccTATTACTTGGGTATTGCCCATAC | Construction of GA pFA6-El222(5xBS)-prSPO13-cln2-mNG |

| | | |
|---|---|---|
| L2-mNG_fwd | ccaagtaataGGTTCTATGGTGAGCAAGGGCGAG | Construction of GA pFA6-El222(5xBS)-prSPO13-cln2-mNG |
| L2-mNG_rev | ccaatacgcaaaccgcctctTTACTTGTACAGCTCGTCCATGC | Construction of GA pFA6-El222(5xBS)-prSPO13-cln2-mNG |
| mNG-rev(cust) | CCAATACGCAAACCGCCTCT | Construction of pFA6-El222(5xBS)-prSPO13-KOZAK-cln2-mNG |
| EL222-spo13-fwd(c) | CATCAGAGCAGATTGTAGAAG | Construction of pFA6-El222(5xBS)-prSPO13-KOZAK-cln2-mNG |
| EL222-spo13-rev(c) | CATTTTGTTttaattaaAATTATTCTCGACTCAACTTC | Construction of pFA6-El222(5xBS)-prSPO13-KOZAK-cln2-mNG |
| cln2-mNG_fwd(auto) | AATTttaattaaAACAAAATGGCTAGTGCTGAACCAAG | Construction of pFA6-El222(5xBS)-prSPO13-KOZAK-cln2-mNG |
| GAL1-UAS_fwd | CGGCATCAGAGCAGATTGTAGAATTTTCAAAAATTCTTACTTTTTTTTTG | Construction of pFA6-GAL1(UP)-EL222(5xBS)-prGAL1-KOZAK-cln2-mNG |
| El222_rev | GAGGACGCACGGCTCTAGTGTCTAAGCTTCATGGACTAAAGGC | Construction of pFA6-GAL1(UP)-EL222(5xBS)-prGAL1-KOZAK-cln2-mNG |
| El222_fwd2 | ATATTGAAGTGGGAGATCTTcGCTAGCCTC | Construction of pFA6-GAL1(UP)-EL222(5xBS)-prGAL1-KOZAK-cln2-mNG |
| GAL1-UAS_rev2 | GAGGCTAGCGAAGATCTCCCACTTCAATATAGCAATGAGC | Construction of pFA6-GAL1(UP)-EL222(5xBS)-prGAL1-KOZAK-cln2-mNG |
| GAL1_fwd | GACACTAGAGCCGTGCGTCCTCGTCTTC | Construction of pFA6-GAL1(UP)-EL222(5xBS)-prGAL1-KOZAK-cln2-mNG |
| GAL1-rev(New) | GCCATTTTGTTTTAATTAATATAGTTTTTTCTCCTTGACGTTAAAGTATAGAGG | Construction of pFA6-GAL1(UP)-EL222(5xBS)-prGAL1-KOZAK-cln2-mNG |
| cln2-fwd-Gal(New) | CTATATTAATTAAAACAAAATGGCTAGTGCTGAACCAAG | Construction of pFA6-GAL1(UP)-EL222(5xBS)-prGAL1-KOZAK-cln2-mNG |

| | | |
|---|---|---|
| mNG_rev_gal | CCAATACGCAAACCGCCTCTTTACTTGTACAGCTCGTCCATGCC | Construction of pFA6-GAL1(UP)-EL222(5xBS)-prGAL1-KOZAK-cln2-mNG |
| Backbone_fwd | attaattaaaacaaaatggctagtg | Construction of pFA6-GAL1(UP)-EL222(5xBS)-prGAL1-KOZAK-cln2-mNG |
| Backbone_rev | aagatctcccacttcaatatagcaatgagc | Construction of pFA6-GAL1(UP)-EL222(5xBS)-prGAL1-KOZAK-cln2-mNG |
| MX terminator rev | cagtatagcgaccagcattc | pACT1-EL222-AQTrip-CYC1t linearization |
| Seq4_fwd | AATTATCCTGGGCACGAG | pACT1-EL222-AQTrip-CYC1t linearization |
| mScarlet_fwd | atatcacaggggccactccactcacGTGAGCAAGGGCGAGGCAGT | Construction of pFA6-Sfp1-mScarlet-NATMX |
| mScarlet_rev | tactaacgccgccatccagtgtcgaTTATTTGTACAGCTCGTCCATGCCG | Construction of pFA6-Sfp1-mScarlet-NATMX |
| pFA6-sfp1-NatMX_fwd_2 | CGAGCTGTACAAATAAtcgacactggatggcggcgtta | Construction of pFA6-Sfp1-mScarlet-NATMX |
| pFA6-sfp1-NATMX-rev_2 | CTCGCCCTTGCTCACGTGAGTGGAGTGGCCCCTGTGATA | Construction of pFA6-Sfp1-mScarlet-NATMX |

**Supplementary Table 4:** List of target sequences and primers for repair fragments used in the study.

| Primer | Target sequence and repair fragments | Source |
|---|---|---|
| cln2 prom target sequence | AACTATTATGCTCCTCTTAC | This study |
| Cln3 cds target sequence | TGCATTAGCGTATCTAATTA | |
| cln1 C-terminal target sequence | GTCACAGTTGAGAGCTATTG | This study |
| gRNA1_fwd_cln1 | GACTTTGTCACAGTTGAGAGCTATTG | This study |
| gRNA1_rev_cln1 | AAACCAATAGCTCTCAACTGTGACAA | This study |
| RF_AQ_EL222_fwd | CCGTTTAGTTCCAACGTAAAATTGTGCCTTTGGACTTAAAATGGCGTgaagcgggtaagctgccac | This study |
| RF_AQ_EL222_rev | ccatgtcgctggccgggtgacccggcggggacgaggcaagctaaacagatctgcaaattaaagccttcgagcg | This study |
| RF_cln2-tail1-fwd | TTGTTTTCTCTACGAGTGAAAGGCATTTTTTGAAACCACTATTCGTTTCCgaagcttGGGAGATCTTCG | This study |
| RF_cln2-tail2-fwd | ACTGTTGGCCCTTTTTGAGGATCTAACCTGCGAAATGTTGATCTGACAGGgaagcttGGGAGATCTTCG | This study |
| RF_cln2-tail-rev | CTCTTTTTGGTACGTTTGGCAAATTGGCATTCATTTATCATGAAAAGAACAGGAATTACTTGTACAGCTCGTCCATGC | This study |

| | | |
|---|---|---|
| Gal_cln2-tail1-f | TTGTTTTCTCTACGAGTGAAAGGCATTTTTTGAAACCACTATTCGTTTCCATGGCTAGTGCTGAACCAAG | This study |
| Gal_cln2-tail2-f | ACTGTTGGCCCTTTTTGAGGATCTAACCTGCGAAATGTTGATCTGACAGGATGGCTAGTGCTGAACCAAG | This study |
| Gal_cln2-tail-rev | CTCTTTTTGGTACGTTTGGCAAATTGGCATTCATTTATCATGAAAAGAACAGGAATTACTTGTACAGCTCGTCC | This study |
| RF_cln1Δ_for | CAATTAAATAAAATAGCACTACCACCACTCCACTGCTCGTTAGCTATTTCTGCCATTTTCTCATCAAGCCTAC | This study |
| RF_cln1Δ_rev | GTAGTATTCCGTTATTAATTAAGTATATATGTAGGCTTGATGAGAAAATGGCAGAAATAGCTAACGAGCAG | This study |
| RF_Whi5_untag_fwd | GCAACAACACCGACCCGTGCCGGCGACATCTGCTGGAGAACCCACGGACGAAACGGAGCCAGAGTC | This study |
| RF_Whi5_untag_rev | CGCGGCTGCACTAACTCCGAGATTGCGGAGAAAAAACTCGTACTACCACATTAAGACGTCTCCACTTCGGTATCCGACTCTGGCTCCGTTTC | This study |
| RF7_cln2_fwd_t1 | TTGTTTTCTCTACGAGTGAAAGGCATTTTTTGAAACCACTATTCGTTTCCGAATTTTCAAAAATTCTTACTTTTTTTTTGG | This study |
| RF7_cln2_fwd_t2 | ACTGTTGGCCCTTTTTGAGGATCTAACCTGCGAAATGTTGATCTGACAGGGAATTTTCAAAAATTCTTACTTTTTTTTTGG | This study |
| RF7_cln2_rev | GAAGTGAGAAAGTAATTCTGCATTAG | This study |

**Supplementary Table 5:** Volumes of daughter cells at birth and budding in G1 cyclin mutant strains.

| Genotype | Volume at birth (median) | Volume at budding (median) | N of cell cycles |
|---|---|---|---|
| CLN1 CLN2 CLN3 | 25.005 fl | 38.755 fl | 82 |
| CLN1 *cln2Δ cln3Δ* | 85.07 fl | 129.985 fl | 44 |
| CLN1 CLN2 *cln3Δ* | 30.62 fl | 52.45 fl | 77 |
| OPTO-CLN2 CLN1 *cln3Δ* Light | 26.18 fl | 33.705 fl | 106 |
| OPTO-CLN2 CLN1 *cln3Δ* Dark | 73.23 fl | 145.215 fl | 100 |
| OPTO-CLN2 *cln1Δ cln3Δ* Light | 18.3 fL | 24.1 fL | 48 |

**Supplementary Table 6:** G1 and post-G1 duration of daughter cells of G1 cyclin mutants.

| Genotype | G1 duration (median) | Post-G1 duration (median) | N of cell cycles |
|---|---|---|---|

| | | | |
|---|---|---|---|
| CLN1 CLN2 CLN3 | 45.0 | 65.0 | 109 |
| CLN1 *cln2Δ cln3Δ* | 100.0 | 55.0 | 79 |
| CLN1 CLN2 *cln3Δ* | 45.0 | 50.0 | 97 |
| OPTO-CLN2 CLN1 *cln3Δ* Light | 35.0 | 70.0 | 106 |
| OPTO-CLN2 CLN1 *cln3Δ* Dark | 100.0 | 60.0 | 61 |
| OPTO-CLN2 *cln1Δ cln3Δ* Light | 40.0 | 75.0 | 72 |

**Supplementary Table 7:** Volumes of mother cells at budding in G1 cyclin mutant strains.

| Genotype | Volume at budding (median) | N of cell cycles |
|---|---|---|
| CLN1 CLN2 CLN3 | 52.585 fl | 70 |
| CLN1 *cln2Δ cln3Δ* | 201.97 fl | 46 |
| CLN1 CLN2 *cln3Δ* | 76.085 fl | 76 |
| OPTO-CLN2 CLN1 *cln3Δ* Light | 40.12 fl | 75 |
| OPTO-CLN2 CLN1 *cln3Δ* Dark | 169.83 fl | 63 |
| OPTO-CLN2 *cln1Δ cln3Δ* Light | 39.07 fl | 38 |

**Supplementary Table 8:** G1 and post-G1 duration of daughter cells of G1 cyclin mutants.

| Genotype | G1 duration (median) | Post-G1 duration (median) | N of cell cycles |
|---|---|---|---|
| CLN1 CLN2 CLN3 | 20.0 | 65.0 | 97 |
| CLN1 *cln2Δ cln3Δ* | 25.0 | 55.0 | 75 |
| CLN1 CLN2 *cln3Δ* | 25.0 | 45.0 | 71 |
| OPTO-CLN2 CLN1 *cln3Δ* Light | 25.0 | 65.0 | 75 |
| OPTO-CLN2 CLN1 *cln3Δ* Dark | 15.0 | 55.0 | 51 |
| OPTO-CLN2 *cln1Δ cln3Δ* Light | 25.0 | 65.0 | 57 |

**Supplementary Table 9:** Statistical comparisons of daughter cell size (Mann-Whitney test p-value, effect size given by rank-biserial correlation).

| Strain comparisons | *p*-value | Effect size (r) | Sample size |
|---|---|---|---|
| CLN1 CLN2 CLN3 - CLN1 CLN2 *cln3Δ* (birth) | 2.89E-05 | 0.38 | 82 / 77 |
| CLN1 CLN2 CLN3 - CLN1 CLN2 *cln3Δ* (bud) | 1.06E-11 | 0.62 | 82 / 77 |
| CLN1 CLN2 CLN3 - CLN1 *cln2Δ cln3Δ* (birth) | 3.14E-20 | 1.0 | 82 / 44 |
| CLN1 CLN2 CLN3 - CLN1 *cln2Δ cln3Δ* (bud) | 2.72E-20 | 1.0 | 82 / 44 |
| CLN1 CLN2 *cln3Δ* - CLN1 *cln2Δ cln3Δ* (birth) | 4.44E-19 | 0.98 | 77 / 44 |
| CLN1 CLN2 *cln3Δ* - CLN1 *cln2Δ cln3Δ* (bud) | 8.34E-20 | 1.0 | 77 / 44 |
| OPTO-CLN2 CLN1 cln3Δ Light – OPTO-CLN2 CLN1 cln3Δ Dark (birth) | 1.75E-35 | 0.99 | 63 / 104 |
| OPTO-CLN2 CLN1 cln3Δ Light – OPTO-CLN2 CLN1 cln3Δ Dark (bud) | 5.92E-36 | 1.0 | 63 / 104 |
| OPTO-CLN2 CLN1 cln3Δ Light – CLN1 CLN2 CLN3 (birth) | 0.99 | 0.0007 | 63 / 82 |
| OPTO-CLN2 CLN1 cln3Δ Light – CLN1 CLN2 CLN3 (bud) | 0.0004 | 0.30 | 63 / 82 |
| OPTO-CLN2 CLN1 cln3Δ Dark - CLN1 cln2Δ cln3Δ (birth) | 0.0008 | 0.35 | 104 / 44 |
| OPTO-CLN2 CLN1 cln3Δ Dark - CLN1 cln2Δ cln3Δ (bud) | 0.30 | -0.11 | 104 / 44 |
| OPTO-CLN2 *cln1Δ* cln3Δ Light – CLN1 CLN2 CLN3 (birth) | 4.44E-10 | 0.66 | 48 / 82 |
| OPTO-CLN2 *cln1Δ* cln3Δ Light – CLN1 CLN2 CLN3 (bud) | 4.52E-14 | 0.79 | 48 / 82 |

**Supplementary Table 10:** Statistical comparisons of daughter G1 and post-G1 durations (Mann-Whitney test p-value, effect size given by rank-biserial correlation).

| Strain comparisons | *p*-value | Effect size (r) | Sample size |
|---|---|---|---|
| CLN1 CLN2 CLN3 - CLN1 CLN2 *cln3Δ* (G1) | 0.92 | 0.0083 | 109 / 97 |
| CLN1 CLN2 CLN3 - CLN1 CLN2 *cln3Δ* (post-G1) | 6.95E-29 | -0.89 | 109 / 97 |
| CLN1 CLN2 CLN3 - CLN1 *cln2Δ cln3Δ* (G1) | 2.73E-26 | 0.91 | 109 / 79 |
| CLN1 CLN2 CLN3 - CLN1 *cln2Δ cln3Δ* (post-G1) | 7.21E-21 | -0.79 | 109 / 79 |

| | | | |
|---|---|---|---|
| CLN1 CLN2 *cln3Δ* - CLN1 *cln2Δ cln3Δ* (G1) | 5.19E-17 | 0.73 | 97 / 79 |
| CLN1 CLN2 *cln3Δ* - CLN1 *cln2Δ cln3Δ* (post-G1) | 3.51E-10 | 0.54 | 97 / 79 |
| OPTO-CLN2 CLN1 cln3Δ Light – CLN1 CLN2 CLN3 (G1) | 1.4E-09 | 0.48 | 106 / 109 |
| OPTO-CLN2 CLN1 cln3Δ Light – CLN1 CLN2 CLN3 (post-G1) | 0.005 | -0.22 | 106 / 109 |
| OPTO-CLN2 CLN1 cln3Δ Light – OPTO-CLN2 CLN1 cln3Δ Dark (G1) | 8.1E-14 | 0.69 | 106 / 61 |
| OPTO-CLN2 CLN1 cln3Δ Light – OPTO-CLN2 CLN1 cln3Δ Dark (post-G1) | 2.84E-12 | -0.64 | 106 / 61 |
| OPTO-CLN2 CLN1 cln3Δ Dark - CLN1 cln2Δ cln3Δ (G1) | 0.24 | 0.12 | 61 / 79 |
| OPTO-CLN2 CLN1 cln3Δ Dark - CLN1 cln2Δ cln3Δ (post-G1) | 0.0001 | -0.37 | 61 / 79 |
| OPTO-CLN2 *cln1Δ* cln3Δ Light – CLN1 CLN2 CLN3 (G1) | 0.12 | 0.13 | 72 / 109 |
| OPTO-CLN2 *cln1Δ* cln3Δ Light – CLN1 CLN2 CLN3 (post-G1) | 6.89E-06 | -0.39 | 72 / 109 |
| OPTO-CLN2 *cln1Δ* cln3Δ Light – OPTO-CLN2 CLN1 cln3Δ Light (G1) | 0.0009 | -0.29 | 72 / 106 |
| OPTO-CLN2 *cln1Δ* cln3Δ Light – OPTO-CLN2 CLN1 cln3Δ Light (post-G1) | 0.09 | 0.12 | 72 / 106 |

**Supplementary Table 11:** Statistical comparisons of mother cell sizes (Mann-Whitney test p-value, effect size given by rank-biserial correlation).

| **Strain comparisons** | ***p*-value** | **Effect size (r)** | **Sample size** |
|---|---|---|---|
| CLN1 CLN2 CLN3 - CLN1 CLN2 *cln3Δ* | 1.03E-18 | 0.85 | 70 / 76 |
| CLN1 CLN2 CLN3 - CLN1 *cln2Δ cln3Δ* | 1.05E-19 | 1.0 | 70 / 46 |
| CLN1 CLN2 *cln3Δ* - CLN1 *cln2Δ cln3Δ* | 2.07E-19 | 0.98 | 76 / 46 |
| OPTO-CLN2 CLN1 cln3Δ Light – CLN1 CLN2 CLN3 | 6.21E-12 | -0.66 | 70 / 75 |
| OPTO-CLN2 CLN1 cln3Δ Light – OPTO-CLN2 CLN1 cln3Δ Dark | 5.74E-24 | 1.0 | 70 / 63 |
| OPTO-CLN2 CLN1 cln3Δ Dark - CLN1 cln2Δ cln3Δ | 0.003 | 0.33 | 63 / 46 |
| OPTO-CLN2 *cln1Δ* cln3Δ Light – CLN1 CLN2 CLN3 | 1.32E-08 | 0.66 | 38 / 70 |

**Supplementary Table 12:** Statistical comparisons of mother G1 and post-G1 durations (Mann-Whitney test p-value, effect size given by rank-biserial correlation).

| Strain comparisons | *p*-value | Effect size (r) | Sample size |
|---|---|---|---|
| CLN1 CLN2 CLN3 - CLN1 CLN2 *cln3Δ* (G1) | 0.2 | 0.11 | 97 / 71 |
| CLN1 CLN2 CLN3 - CLN1 CLN2 *cln3Δ* (post-G1) | 2.28E-20 | -0.83 | 97 / 71 |
| CLN1 CLN2 CLN3 - CLN1 *cln2Δ cln3Δ* (G1) | 0.023 | 0.2 | 97 / 75 |
| CLN1 CLN2 CLN3 - CLN1 *cln2Δ cln3Δ* (post-G1) | 6.42E-09 | -0.51 | 97 / 75 |
| CLN1 CLN2 *cln3Δ* - CLN1 *cln2Δ cln3Δ* (G1) | 0.38 | 0.083 | 71 / 75 |
| CLN1 CLN2 *cln3Δ* - CLN1 *cln2Δ cln3Δ* (post-G1) | 6.31E-08 | 0.51 | 71 / 75 |
| OPTO-CLN2 CLN1 cln3Δ Light – CLN1 CLN2 CLN3(G1) | 0.001 | -0.28 | 75 / 97 |
| OPTO-CLN2 CLN1 cln3Δ Light – CLN1 CLN2 CLN3 (post-G1) | 0.4 | -0.07 | 75 / 97 |
| OPTO-CLN2 CLN1 cln3Δ Light – OPTO-CLN2 CLN1 cln3Δ Dark (G1) | 4.5E-07 | -0.52 | 75 / 51 |
| OPTO-CLN2 CLN1 cln3Δ Light – OPTO-CLN2 CLN1 cln3Δ Dark (post-G1) | 7.06E-08 | -0.55 | 75 / 51 |
| OPTO-CLN2 CLN1 cln3Δ Dark - CLN1 cln2Δ cln3Δ (G1) | 5.67E-06 | 0.47 | 51 / 75 |
| OPTO-CLN2 CLN1 cln3Δ Dark - CLN1 cln2Δ cln3Δ (post-G1) | 0.53 | -0.06 | 51 / 75 |
| OPTO-CLN2 *cln1Δ* cln3Δ Light – CLN1 CLN2 CLN3 (G1) | 0.04 | -0.19 | 57 / 97 |
| OPTO-CLN2 *cln1Δ* cln3Δ Light – CLN1 CLN2 CLN3 (post-G1) | 0.008 | -0.25 | 57 / 97 |
| OPTO-CLN2 *cln1Δ* cln3Δ Light – OPTO-CLN2 CLN1 cln3Δ Light (G1) | 0.85 | 0.02 | 57 / 75 |
| OPTO-CLN2 *cln1Δ* cln3Δ Light – OPTO-CLN2 CLN1 cln3Δ Light (post-G1) | 0.09 | -0.17 | 57 / 75 |

**Supplementary Table 13:** Statistical comparisons of daughter specific volume increase during G1 (Mann-Whitney test p-value, effect size given by rank-biserial correlation.

| Strain comparisons | *p*-value | Effect size (r) | Sample size |
|---|---|---|---|
| CLN1 CLN2 CLN3 – CLN1 cln2Δ cln3Δ | 0,37 | 0,1 | 82 / 44 |
| CLN1 CLN2 CLN3 – OPTO - CLN2 CLN1 cln3Δ Light | 0,06 | 0,18 | 82 / 57 |

| | | | |
|---|---|---|---|
| OPTO-CLN2 CLN1 cln3Δ Dark - OPTO-CLN2 CLN1 cln3Δ Light | 0,08 | -0,17 | 103 / 57 |
| OPTO-CLN2 CLN1 cln3Δ Dark - CLN1 cln2Δ cln3Δ | 0,02 | -0,33 | 103 / 44 |
| OPTO-CLN2 CLN1 cln3Δ Dark - CLN1 CLN2 CLN3 | 3.89E-06 | -0,39 | 103 / 82 |
| OPTO-CLN2 CLN1 cln3Δ Light - CLN1 cln2Δ cln3Δ | 0,31 | -0,12 | 57 / 44 |
| CLN1 CLN2 CLN3 - OPTO-CLN2 *cln1 cln3Δ* arrest | 0.001 | 0.32 | 82 / 64 |
| OPTO-CLN2 *cln1 cln3Δ* Light - OPTO-CLN2 *cln1Δ cln3Δ* arrest | 0.19 | -0.15 | 47 / 64 |
| CLN1 CLN2 CLN3 - OPTO-CLN2 *cln1Δ cln3Δ* Light | 0.004 | 0.3 | 82 / 47 |
